# AMH protects the ovary from doxorubicin by regulating cell fate and the response to DNA damage

**DOI:** 10.1101/2024.05.23.595356

**Authors:** Ngoc Minh Phuong Nguyen, Eun Mi Chang, Maeva Chauvin, Natalie Sicher, Aki Kashiwagi, Nicholas Nagykery, Christina Chow, Phoebe May, Alana Mermin-Bunnel, Josephine Cleverdon, Thy Duong, Marie-Charlotte Meinsohn, Dadi Gao, Patricia K. Donahoe, David Pepin

**Affiliations:** Pediatric Surgical Research Laboratories, Massachusetts General Hospital, MA, USA; Department of Surgery, Harvard Medical School, Boston, MA, USA; Department of Neurology, Department of Neurology, Massachusetts General Hospital and Harvard Medical School; Center for Genomic Medicine, Massachusetts General Hospital Research Institute, Boston, MA

## Abstract

Anti-Müllerian hormone (AMH) protects the ovarian reserve from chemotherapy, and this effect is most pronounced with Doxorubicin (DOX). However, the mechanisms of DOX toxicity and AMH rescue in the ovary remain unclear. Herein, we characterize these mechanisms in various ovarian cell types using scRNAseq. In the mesenchyme, DOX activates the intrinsic apoptotic signaling pathway through p53 class mediators, particularly affecting theca progenitors, while co-treament with AMH halts theca differentiation and reduces apoptotic gene expression. In preantral granulosa cells, DOX upregulates the cell cycle inhibitor *Cdkn1a* and dysregulates Wnt signaling, which are ameliorated by AMH co-treatment. Finally, in follicles, AMH induces *Id3*, a protein involved in DNA repair, which is necessary to prevent the accumulation of DNA lesions marked by γ-H2AX in granulosa cells. Altogether this study characterizes cell, and follicle stage-specific mechanisms of AMH protection of the ovary, offering promising new avenues for fertility preservation in cancer patients undergoing chemotherapy.

**Highlights:**

- Doxorubicin treatment induces DNA damage that activates the p53 pathway in stromal and follicular cells of the ovary.
- AMH inhibits the proliferation and differentiation of theca and granulosa cells and promotes follicle survival following Doxorubicin insult.
- AMH treatment mitigates Doxorubicin-induced DNA damage in the ovary by preventing the accumulation of γ-H2AX-positive unresolved foci, through increased expression of ID3, a protein involved in DNA repair.

**Graphical Abstract:** 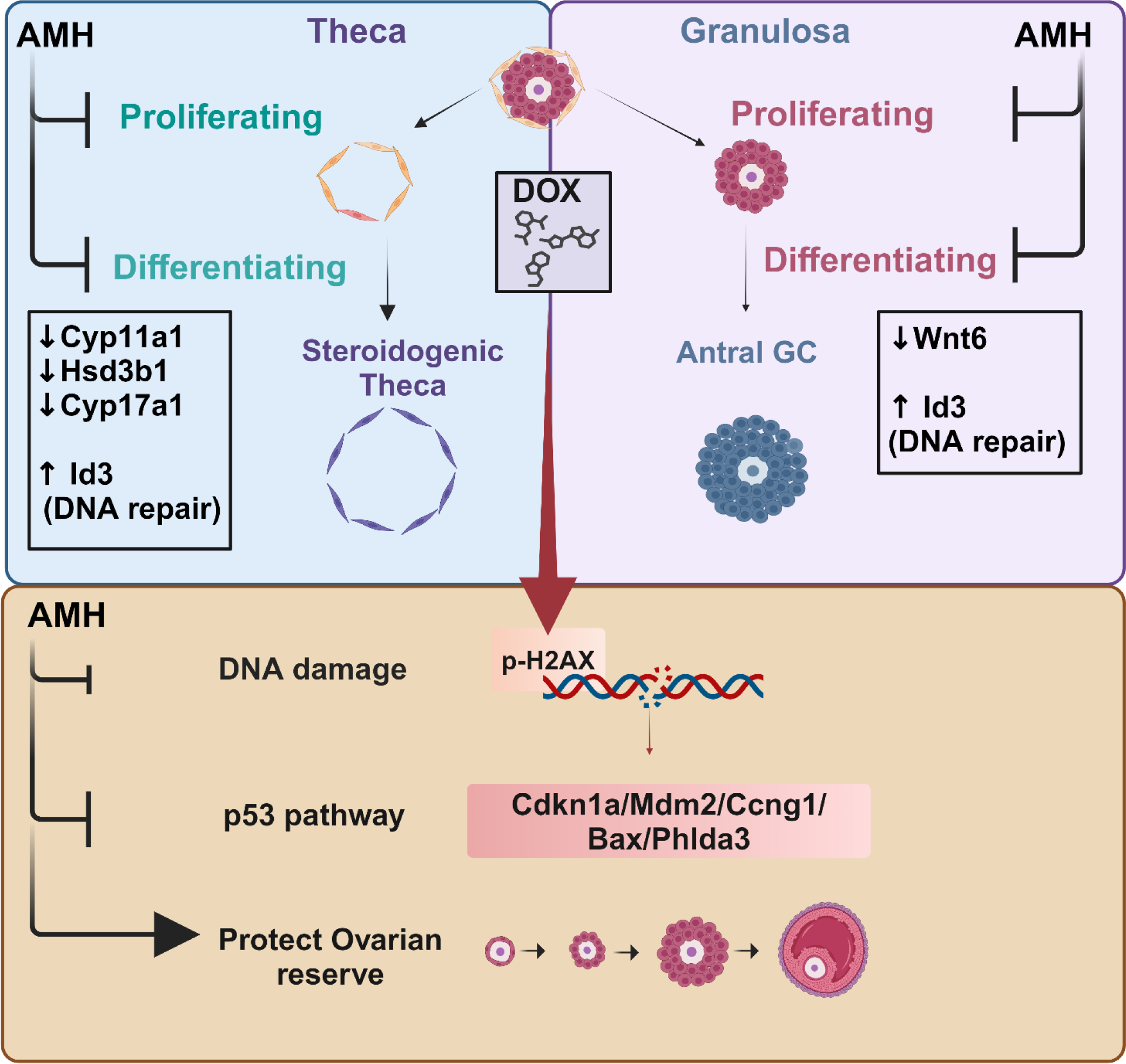

## Introduction

The adverse effects of chemotherapy on future fertility have long been recognized^1,2^. Over the past few decades, survivorship of cancer therapies has improved significantly, leading to an increasing need to understand and prevent the adverse long-term impacts of these treatments on fertility, especially in pediatric patients and young adults. The standard of care for preserving reproductive function is gamete or embryo freezing, and increasingly ovarian cortex tissue cryopreservation before cancer therapy ^3^. However, these methods are not available to all patient populations, can delay cancer treatment, and in the case of tissue transplantation may risk reintroducing cancer cells^3^. For these reasons, there is a clinical need for treatments that preserve fertility (fertoprotectants) that can be given concurrently with chemotherapy; GnRH analogs are often used for this purpose, with the hypothesis that their suppressive effect on antral follicles may be salutary to the ovary, although their clinical efficacy is limited^4–6^ .

One potential fertoprotectant that has been proposed is Anti-Mullerian Hormone (AMH)^7^, a member of the transforming growth factor-beta (TGFβ) superfamily of ligands that is produced by granulosa cells of growing follicles and is a feedback inhibitor of follicle activation and pre- antral follicle growth^8,9^. Currently, measurements of circulating AMH are used in the clinic as an marker of ovarian reserve (the pool of quiescent primordial follicles), and to predict fertility and reproductive outcomes after chemotherapy^10^. In addition to being a biomarker, AMH itself is a potent suppressant of early follicle development : studies have shown that treatment with exogenous AMH can provide durable contraception in female mice, rats, and cats^11–13^. Because AMH suppresses follicles at an earlier stage than GnRH analogs, it has the potential to be more efficacious in safeguarding the much larger pool of preantral follicles and the ovarian reserve from the damages of chemotherapy. Indeed, we and others have shown that AMH may help protect the ovarian reserve during exposure to chemotherapy^11,14–17^. Co-treatment with recombinant human AMH (rhAMH) significantly reduced the loss of primordial follicles in mice treated with carboplatin, cyclophosphamide, and Doxorubicin^11^. One presumed mechanism of action is the replacement of the negative feedback normally provided by endogenous AMH from growing follicles on new follicle activation and growth, thus preventing the excessive activation of follicles that has been speculated to occur when treating with chemotherapy^18^. The protective effect of exogenous AMH was particularly pronounced in conjunction with Doxorubicin (DOX)^11^, an anthracycline chemotherapeutic drug and a topoisomerase inhibitor ^19^ used to treat a wide variety of cancers. DOX disrupts DNA replication and induces DNA lesions, making it especially harmful to dividing cells. In contrast, in our model, AMH was less effective at protecting the ovarian reserve from the damage of cyclophosphamide^11^, which is particularly damaging to mitotically arrested primordial oocytes, suggesting AMH may be most useful in the context of chemotherapies that target dividing granulosa cells of growing follicles^20^.

Considering the promising clinical potential of AMH, and the clinical need for such treatment in pediatric patients, mechanistic studies are necessary to understand the effects of repeated chemotherapy cycles on ovarian follicle reserve and the protective role of AMH against such insults. In this study, we aimed to explore the mechanisms behind ovarian damage in prepubertal mice following DOX treatment, and to elucidate the mechanisms by which AMH may prevent chemotherapy-induced ovarian damage, using single-cell RNA sequencing (scRNA-seq), as well as quantitative and morphometric histological evaluations. We found that DOX induced *Cdkn1a* (P21)-driven DNA-damage pathways in multiple ovarian cell types, and that unexpectedly, AMH may protect the ovary both by inhibiting progenitor proliferation and differentiation of theca and granulosa cells, but also by ameliorating the resolution of DNA damage through the upregulation of *Id3*.

## Results

### Classification of cell types in mouse prepubertal ovaries

To evaluate the response of different cell types to DOX damage and AMH treatment we treated mice (n=5) at the prepubertal stage (postnatal day 25) with two weekly doses of DOX (3mg/kg) (Figure 1A). The use of this dose and repeated treatment with DOX allowed us to explore both the acute and chronic effects of DOX and replicate better the clinical protocols using repeated rounds of chemotherapy in pediatric cancer patients. To evaluate the protective effects of AMH during DOX exposure we co-treated mice with recombinant human AMH (rhAMH) (0.75mg/kg/day), a dose we showed previously to help preserve the ovarian reserve^11^. For controls, we also included a group that received vehicle or rhAMH alone. Ovaries were collected at either 4 hours or 1 week after the second dose of DOX.

**Figure 1:**
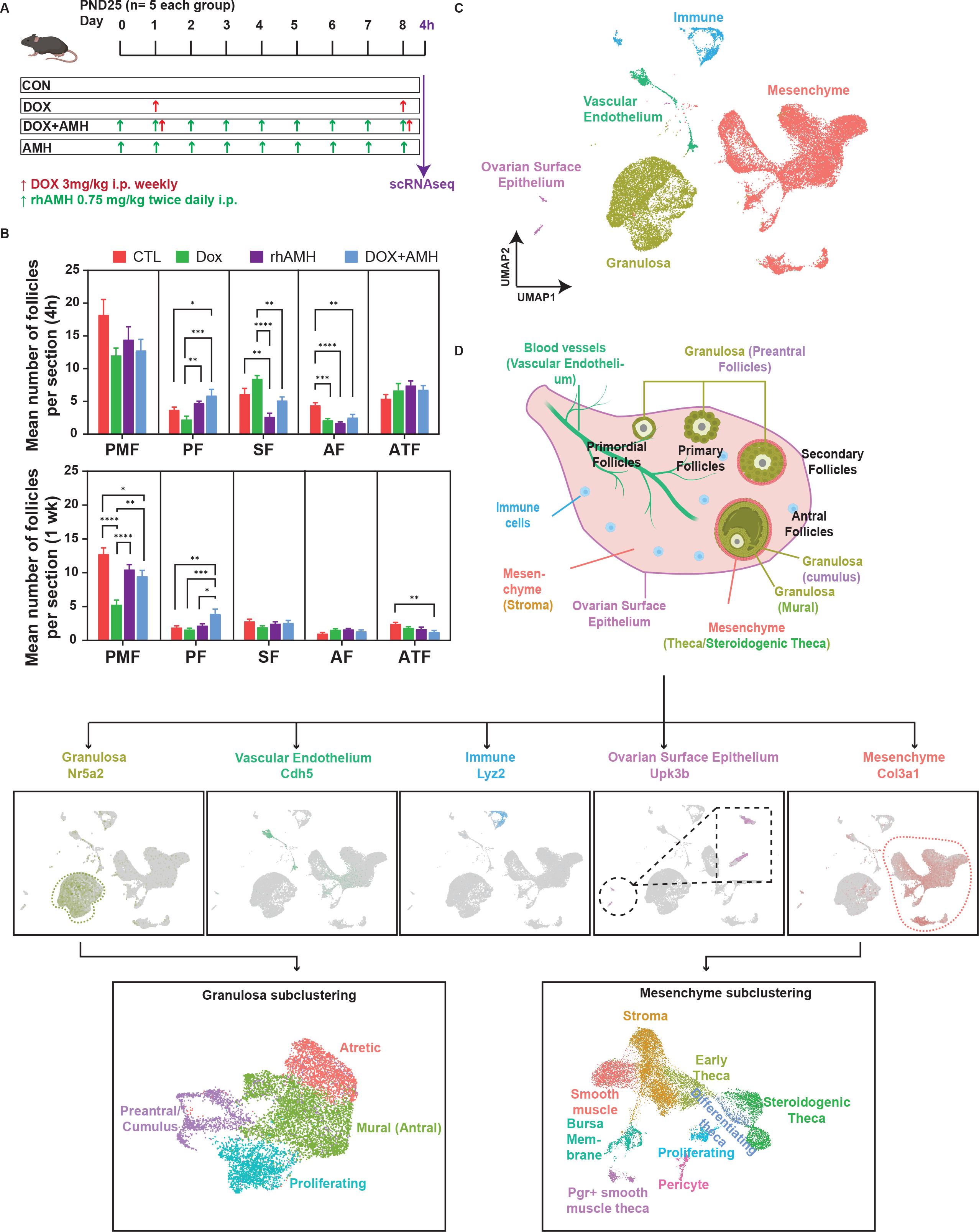
Single-cell RNA sequencing analysis of mouse ovaries treated with DOX and AMH. (A) Mice were treated with DOX and AMH, and ovaries were dissociated into a single-cell suspension for scRNAseq library preparation. (B) The mean number of follicles at different developmental stages in 4 groups at 4hr and 1 week. Data are presented as mean ± SEM; ∗p < 0.05, ∗∗p < 0.01, ∗∗∗p < 0.001. (C) UMAP plot featuring five major cell types in the ovary. (D) Illustration depicting the internal components of the mouse ovary with examples of markers used for identifying five major cell types and the clustering of granulosa cells and mesenchyme.

To examine the protective effect of recombinant rhAMH against DOX-induced ovarian follicle loss^11^, we counted follicles at 4 hours and 1 week after the second DOX dose (Figure 1A). Both AMH alone, or as a co-treatment at 4 hours post-DOX, produced a significant increase in the number of primary ovarian follicles and a significant decrease in secondary and antral follicles (Figure 1B), consistent with the suppressive effect of AMH on preantral follicle development. At the one-week timepoint post-DOX, we observed a substantial reduction in the primordial follicle (PMF) reserve by 59% in DOX-alone treated mice (5.25 ± 0.71/section) compared to controls (12.76 ± 0.92/section, P<0.0001). Importantly, AMH co-treatment significantly mitigated this loss, resulting in an 80% recovery in PMF (9.47 ± 0.87/section in DOX + rhAMH, P< 0.001) compared to DOX-only treated mice (Figure 1B).

To understand more clearly the mechanisms of DOX injury and AMH rescue at the cellular level we used the 10X Genomics workflow, to generate single-cell transcriptomes from ∼10,000 cells in each condition: vehicle control, AMH, DOX, DOX+AMH (Figure 1A). After quality check and normalization, we removed clusters of cells that exhibited stress response genes such as Fos, Fosb, and Jun, or a high proportion of mitochondrial and ribosomal genes counts, likely due to tissue processing and dissociation artifacts. Clusters corresponding to oviductal or rete ovarii cells^21,22^ (expression of *Ovgp1*) and reticulocytes (expression of *Hbb-bs, Hba-a1, Hba-a2, Hbb-bt*) were also excluded from the analysis in this study due to their limited representation. After data cleanup, five major cell types were identified based on previously reported markers^23^: Mesenchyme (58%), Granulosa (GC) (33%), Immune (5%), Vascular Endothelium (3%), Ovarian Surface Epithelium (OSE)(1%) (Figure 1C, Figure S1A). We next re-clustered GC cells and resolved sub-cell types expressing established markers: Preantral/Cumulus (*Gatm*+), Antral-Mural (*Inhba*+), Proliferating (*Top2a*+), and Atretic (*Ghr*+) (Figure 1C, Figure S1B). The reclustering of mesenchymal cells also recapitulated previously reported cell subtypes: smooth muscles (*Myh11*+), Stroma (*Tcf21*+), Proliferating (*Top2a*+), Pericyte (*Rgs5*+), Early Theca (*Hhip*+), Steroidogenic Theca (*Star*+), and Differentiating Theca, which expressed markers of both early theca stage and Steroidogenic Theca (*Hhip+ Star+*) (Figure 1C, Figure S1C). Furthermore, clusters corresponding to cells of the ovarian bursa membrane (*Clec3b*+, validated by RNAish) and *Pgr*+ smooth muscle theca cells (*Pgr*+, validated by RNAish) were also identified (Figure S1C).

### DOX treatment induced expression of p53 targets in mesenchymal cells, which were suppressed by AMH co-treatment

To elucidate the cellular response of mesenchyme cells to DOX, we compared the transcriptomes of the mesenchymal cell clusters between the DOX and CON conditions (Figure 2A, Figure S2A). Differential Gene Expression (DGE) and Gene Ontology (GO) analysis revealed that the most significantly enriched pathway was the intrinsic apoptotic signaling pathway by p53 class mediator, notably *Cdkn1a, Ccng1, Mdm2, Phlda3* and *Bax*, which were also among the top 10 markers that were highly differentially expressed in DOX treatment (Figure 2B, Table S1). GO analysis also revealed a significant upregulation of pathways that regulate muscle cell differentiation (*Wnt4, Lama2, Tmsb4x, Ank2*), mitotic DNA checkpoint signaling (*Ccng1, Cdkn1a, Rps27l, Mdm2*), and smooth muscle cell proliferation (*Cdkn1a/Igf1r/Mdm2/Prkg1*) (Figure 2B, S2A, S2B and Table S2). To investigate further the impact of DOX treatment on proliferating cells, a previously recognized target of DOX^24^, we performed Gene Set Enrichment Analysis (GSEA) on the proliferating mesenchymal cluster using the M2 mouse collection, containing 2626 gene sets from pathway databases and biomedical literature. GSEA revealed significant enrichment in TP53 targets (Ongusaha 2003 dataset)^25^ and in the cellular response of different cell types to DNA-damaging treatments such as Cisplatin, Methotrexate, Camptothecin (Brachat, 2002 dataset)^26^, ionizing radiation (IR) (Quintens dataset)^27^ (Figures 2C, S2C, and Table S3), all sharing common genes involved in the p53 pathway as mechanisms of DNA-damage response. Importantly, co-treatment with AMH resulted in a reduced expression of these p53 target genes as seen in the scRNAseq dataset (Fig 2D-F). Furthermore, we also observed enrichment in the WikiPathways PeroxireDOXin 2- induced ovarian failure pathway (WikiPathways) and ovarian aging (Custom gene sets of aging and senescence signatures extracted from Lengyel 2022^22^) (Figure S2C). Similarly, the ovarian surface epithelial (OSE) cluster recapitulated a signature of p53 target genes upon DOX treatment (Figure S2D and S2E). Altogether, these data suggest that the DOX toxicity to ovarian mesenchymal and ovarian surface epithelial cells elicits a common upregulation of p53 target genes, leading to cell cycle arrest (*Cdkn1a*), Apoptosis (*Bax, Phlda3*), Senescence (*Cdkn1a*), or activation of P53 feedback mechanisms (*Mdm2, Ccng1*) in response to DNA damage and that AMH may dampen these responses (Figure 2G).

**Figure 2.**
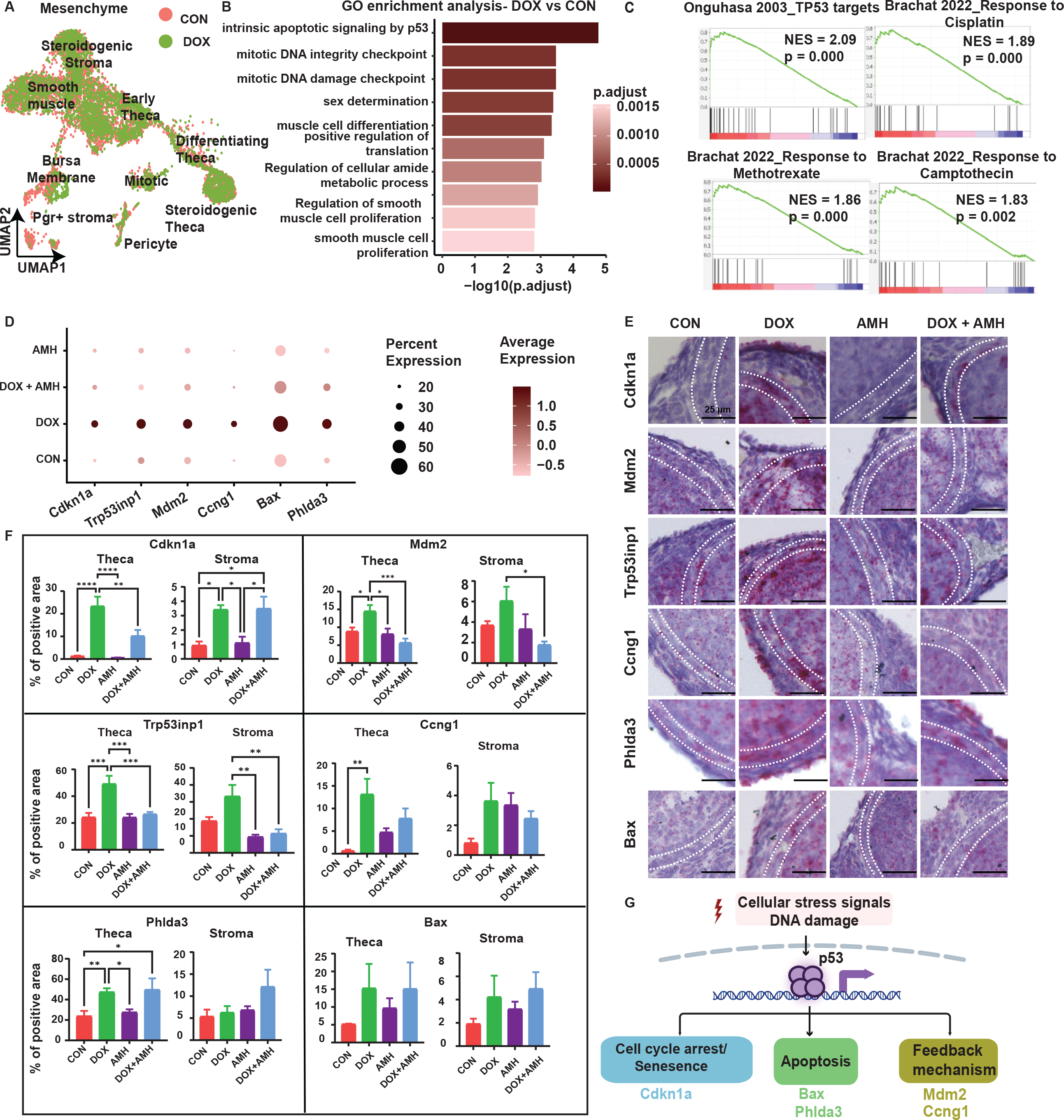
DOX impact on ovarian mesenchymal cells. (A) UMAP of mesenchymal subtypes in DOX-treated and control ovaries. (B) Top 10 GO enriched pathways in response to DOX. (C) Selected GSEA pathways using the M2 collection of 2626 gene sets, highlighting p53 pathway markers in proliferating mesenchymal cells in response to DOX. (D) Expression patterns of the most significantly upregulated 53 pathway markers in response to DOX. (E-F) Representative micrograph of an ovarian section stained by RNAish for selected p53 markers in stroma and theca cells, scale bars = 25 µm (E), along with quantification (F). Data are presented as mean ± SEM; ∗p < 0.05, ∗∗p < 0.01, ∗∗∗p < 0.001. (G) Schematic model of suggested mechanisms activated by DOX.

To validate the upregulations of the p53 target markers in the cell types of interest, we chose to quantify the most prominent genes by fold change of significant markers identified through DGE analysis using RNAish and image analysis (Figure 2F). These markers included *Cdkn1a, Trp53inp1, Mdm2, Ccng1, Phlda3, and Bax*. For each ovary (N= 5), theca layers from 3 growing follicles and 3 random stromal areas per mouse were selected for quantification (Figure S2F). RNA in-situ hybridization (RNAish) quantification confirmed significant upregulation of *Cdkn1a* in both theca and ovarian stroma (Figure 2F). Other p53 target genes (*Trp53inp1*, *Mdm2*, *Ccng1*, *Phlda3*) displayed a significant upregulation in the theca cells, while the ovarian stroma showed a trend towards increased expression, although this increase did not reach statistical significance. Notably, co-treatment with AMH significantly reduced the upregulation of p53 targets (*Cdkn1a*, *Mdm2*, and *Trp53inp1*) in theca cells. Taken together, these data suggest that the upregulation of p53 target genes (such as *Cdkn1a*) in response to DNA damage was prominent in mesenchymal cells following treatment with DOX and that co-treatment with AMH significantly attenuated the induction of these gene targets, particularly in theca cells.

### AMH stalls the proliferation and differentiation of theca cells, contributing to the resistance to DOX

To further investigate the theca cell response to AMH, we first examined the expression of *Amhr2* in prepubertal ovaries. Both the scRNAseq data and the in situ hybridization confirmed the expression of *Amhr2* in granulosa, ovarian surface epithelium, and importantly, in steroidogenic theca cells (Figures S3A and S3B). We focused on the steroidogenic theca cells, which showed a significantly reduced expression of p53 targets when co-treated with AMH compared to DOX alone. Upon AMH treatment, we identified a new theca cluster in the scRNAseq data uniquely associated with AMH treatment (DOX+AMH, AMH) and not other conditions (CON, DOX) (Figures 3A and 3B). All theca clusters (CON, DOX, AMH, DOX+AMH) expressed Prss35, a known serine protease marker expressed in the theca layer of developing follicles^23^ (Figure 3B). DGE analysis contrasting AMH-treated (DOX+AMH, AMH) from those without AMH (CON, DOX) revealed a significant reduction in the expression of key steroidogenic markers such as *Cyp17a1, Cyp11a1,* and *Hsd3b1* in AMH-treated theca cells (Figures 3C and 3D), and GO analysis confirmed enrichment of markers involved in steroid metabolic process (Figure S3C)). Furthermore, *Id3*, an inhibitor of differentiation gene ^28^, and known target of AMH^8^, was found to be upregulated by AMH treatments (Figure 3C). RNAish confirmed the reduced expression of *Cyp11a1* and the near complete absence of *Cyp17a1* expression in theca cells of growing follicles in response to AMH (Figures S3d and 3E). Taken together, these data suggest that AMH treatment stalls or prevents early theca differentiation into steroidogenic theca.

**Figure 3.**
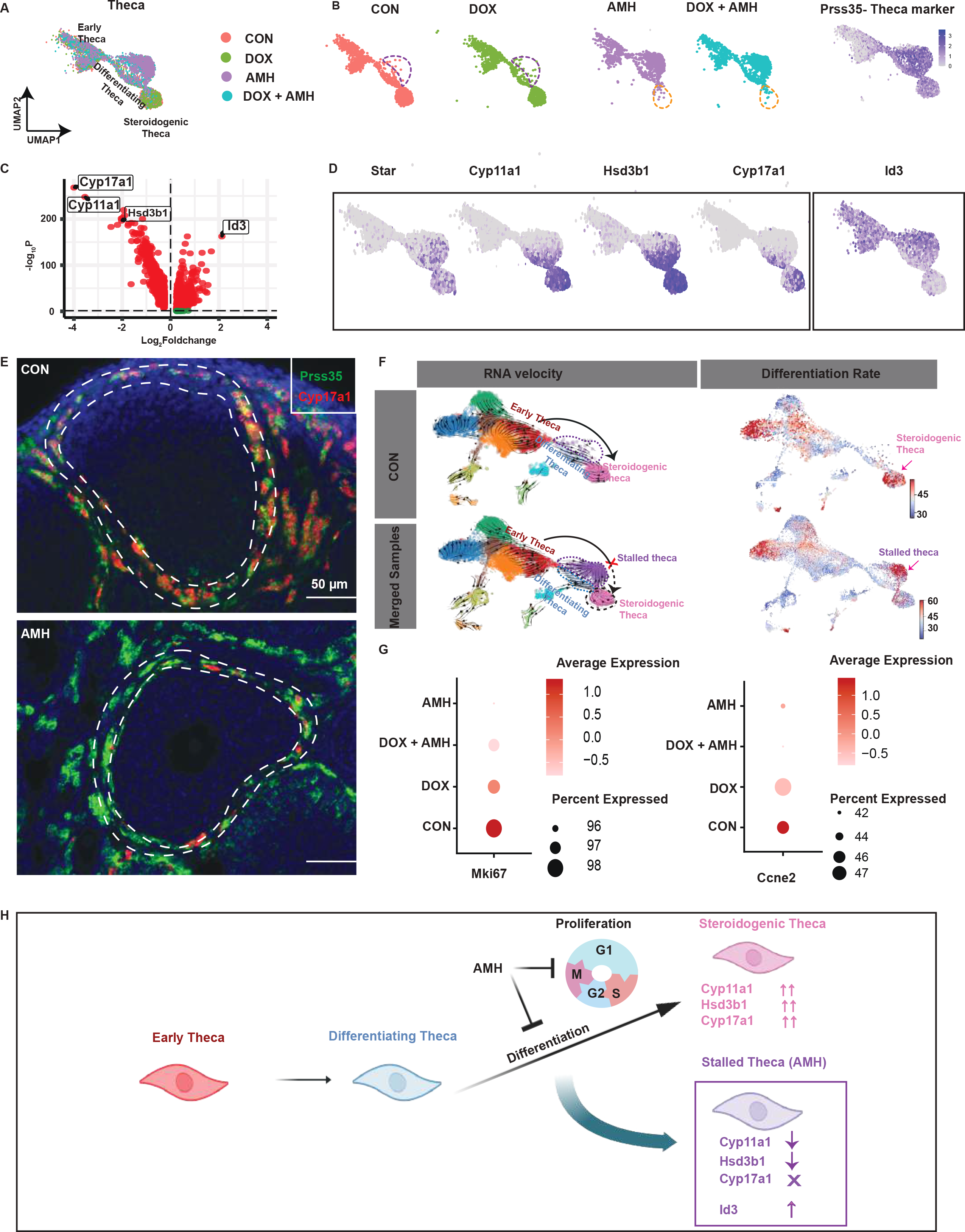
AMH impact on ovarian theca. (A-B) UMAP of theca subclusters (A) under different conditions showing AMH-specific cluster and (B) theca marker *Prss35*. (C) Volcano plot highlighting differentially expressed genes between AMH-treated and control. (D) FeaturePlot showing downregulated steroidogenic markers in AMH-specific clusters and upregulated marker *Id3*. (E) Representative micrograph of an ovarian section stained for *Cyp17a1* by RNAish in control and AMH-treated groups. Scale bars = 50 µm. (F) RNA velocity predicting the differentiation path (left panel) and differentiation rate (right panel) of steroidogenic theca in control and DOX with AMH co-treatment, with arrows pointing to the theca clusters with the highest differentiation rates. (G) Dotplot of proliferation markers *Mki67 and Ccne2* in the proliferating mesenchymal cells. (H) Schematic overview of a suggested model for the differentiation stall induced by AMH.

We then assessed the effects of AMH on cellular differentiation trajectories using RNA velocity analysis. In control ovaries, RNA velocity vectors formed a strong directional flow originating from early theca, passing through differentiating theca, and ending at the terminally differentiated state of steroidogeneic theca, which is marked by high expression of steroidogenic enzymes (Figure 3F, top panel). Differentiation rate analysis by RNA velocity in control ovaries identified this steroidogenic theca state as the terminally differentiated stage in their developmental lineage (Figure 3F, top panel). In contrast, when AMH samples were combined in the RNA velocity analysis, it revealed an AMH-specific cluster consistent with a stall in differentiation as their terminal cell state, instead of terminating at steroidogenic theca (Figure 3F, bottom panel). These findings suggest that AMH alters the cellular differentiation trajectory of progenitor theca cells, causing them to stall in their differentiation process rather than progressing into steroidogenic differentiation.

Given AMH’s role in inhibiting proliferation and preventing cell differentiation in granulosa cells^8^, we next examined the expression of proliferation markers *Mki67* and *Ccne2* in proliferating mesenchymal cells in the scRNASeq dataset. Our analysis confirmed the downregulation of these markers (Figure 3G). We further examined AMH’s antiproliferative effect by quantifying the expression of Ki67, a G2M marker, in theca cells using immunohistochemistry. The proportion of Ki67+ theca cells was lower in DOX than CON, suggesting DOX may eliminate actively dividing cells. While AMH treatment also resulted in a lower percentage of Ki67+ cells at 4 hours, we speculate this was due to temporary inhibition of theca proliferation rather than cell death (Figure S3E). Supporting this interpretation, 1-week after the final AMH treatment, theca cells had a higher rate of proliferation than controls, including in ovaries co-treated with DOX and AMH (Figure S3E).

Altogether, these observations suggest that AMH may temporarily inhibit progenitor theca cell proliferation and stall their differentiation towards the terminal steroidogenic cell state (Figure 3H). This may impart temporary protection of follicles from DOX-induced damage, which would otherwise eliminate proliferating early theca. Following cessation of AMH treatment, theca cells resumed their proliferation and differentiation to recover their steroidogenic functions, which are essential to support follicular development.

### DOX treatment induced p53 targets in preantral granulosa cells, which were rescued by AMH

Given that preantral granulosa cells are highly regulated by AMH, as evidenced by the distinct shifts in clusters of AMH-treated samples by scRNA-seq (Figure 4A), we aimed to understand the impact of DOX on these cells. To elucidate the mechanisms of DOX damage to preantral granulosa cells we performed GSEA analysis, which revealed a significant enrichment of 28 GSEA gene sets (p< 0.05), including notably the upregulation of the p53 target pathways, as seen in the analysis of mesenchymal cells (Table S4). GSEA analysis also revealed a significant upregulation of pathways that regulate cell cycle and DNA damage responses including DNA repair, the p53 pathway, and the Wnt/Beta-Catenin signaling (Figures 4B-D). We validated selected p53 targets by RNAish which confirmed the significant upregulation of *Cdkn1a, Mdm2,* and downstream markers of p53-induced apoptosis Bax (Figure 4E and S4A). Co-treatment with AMH significantly reduced the upregulation of p53 targets *Cdkn1a* and *Mdm2*, and similar trend was observed for the apoptosis marker Bax but did not reach statistical significance (Figure 4E, Figure S4A). Taken together, these findings suggest that DOX may activate the p53 pathway, resulting in the upregulation of *Cdkn1a* and *Mdm2*, and that AMH co-treatment may dampen this p53 response.

**Figure 4.**
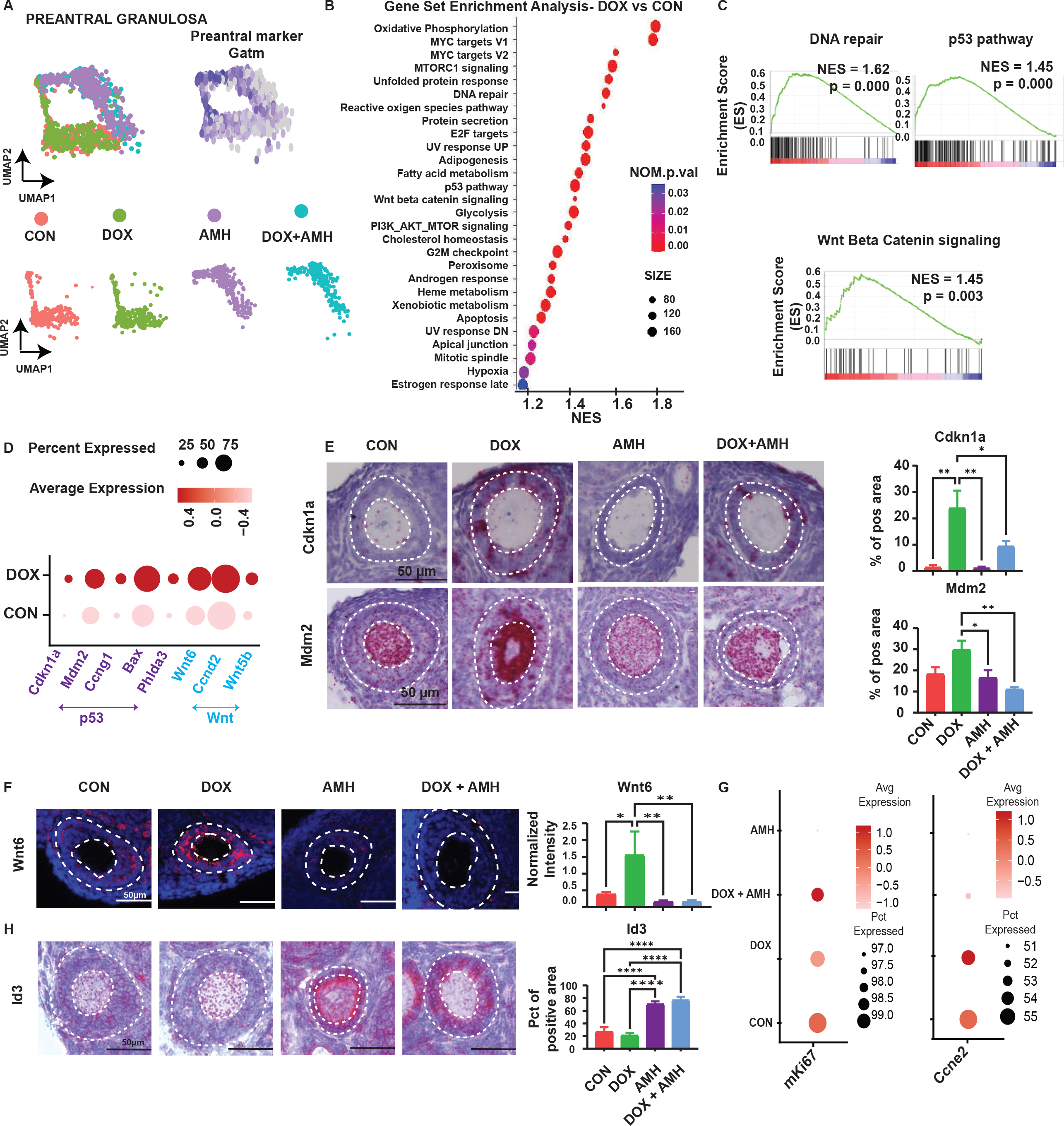
DOX impact on ovarian preantral granulosa cells. (A) UMAP of preantral clusters identified by canonical marker and subtypes in all conditions. (B) All 28 GSEA-enriched pathways in response to DOX. (C) Selected GSEA plots demonstrating pathways that regulate the DNA damage and Wnt Beta Catenin signaling. (D) Expression patterns of markers involved in the p53 and Wnt pathways. (E) Representative micrograph of ovarian section stained by RNAish for selected p53 markers in preantral granulosa, with quantification. Scale bars, 50 µm. Data are presented as mean ± SEM; ∗p < 0.05, ∗∗p < 0.01, ∗∗∗p < 0.001. (F) Representative micrograph of an ovarian section stained by fluorescent RNAish for *Wnt6* showing expression in preantral GC with quantification. Scale bars, 50 µm. Data are presented as mean ± SEM; ∗p < 0.05, ∗∗p < 0.01, ∗∗∗p < 0.001. (G) Dotplot of proliferation markers, *Mki67 and Ccne2*, in proliferating granulosa cells. (H) Representative micrograph of ovarian section stained by RNAish for *Id3* in preantral granulosa, along with quantification. Data are presented as mean ± SEM; ∗p < 0.05, ∗∗p < 0.01, ∗∗∗p < 0.001.

### Doxorubicin treatment induced the expression of Wnt6, while AMH treatment inhibited *Wnt6* and induced *Id3* expression

To explore the effect of DOX on cell-cell communication in preantral follicles we employed CellChat, a computational tool that analyzed and compare intercellular communication network ^29^, and identified signaling pathways altered by DOX. We compared the strength of outgoing and incoming interactions to and from preantral follicle granulosa cells (Figure S4B). Among the enhanced outgoing interactions, the Wnt signaling pathway emerged as the most prominent, indicating that under DOX conditions, preantral follicles upregulated paracrine Wnt signals to other cells, effectively dysregulating this pathway. Given the identification of Wnt signaling as altered by DOX treatment in both GSEA and CellChat analyses upon DOX treatment, and the significant upregulation of *Wnt6* by DOX in the DGE analysis, we validated this change of expression by RNAish and quantified its normalized mean intensity across different treatment conditions (Figure 4F). Increased *Wnt6* expression was observed in primordial, primary, and secondary follicles (collectively considered as pre-antral stages) in DOX-treated ovaries compared to controls. Conversely, *Wnt6* expression was nearly absent in pre-antral follicles treated with either AMH or DOX+AMH (Figure 4F). The contrasting expression patterns of Wnt6 in DOX and AMH-treated samples suggest an antagonism of DOX and AMH on follicle differentiation. In support of this hypothesis, we found inhibition of preantral markers *Slc18a1, Nr5a2, Aldh1a1,* and *Kctd14* (Figs4C) and inhibition of proliferation in granulosa cells with AMH treatment (Figure 4G) in our scRNAseq dataset, as well as by qPCR in whole ovaries (Figure S4D). Thus, we speculate that AMH may protect preantral follicles by inhibiting their proliferation and differentiation which decreases their vulnerability to DOX cytotoxicity.

To understand the response of preantral granulosa cells to AMH, we examined the expression of downstream canonical targets of AMH in granulosa cells^8^. The scRNA-seq analysis confirmed the presence of an AMH-specific preantral cluster gene signature as previously identified^8^ (Figure S4C). In particular, we found a significant increase in *Id3* expression, a protein involved not only in inhibition of differentiation but also in DNA repair and recruited to sites of DNA damage to facilitate double-strand break repair through its interaction with MDC1^28,30^. Using RNAish we confirmed a significant upregulation of *Id3* in AMH-treated preantral granulosa cells (Figure 4H). Interestingly, *Id3* was also strongly induced in the oocytes of AMH-treated preantral follicles (Figure S4E). We speculate that the upregulation of *Id3* in the oocytes and granulosa cells of preantral follicles by AMH may facilitate DNA damage repair following DOX treatment.

### Co-treatment with AMH dampened DOX-induced DNA damage response, and enhanced DNA repair in an Id3-dependent fashion

Next, we wanted to elucidate how DOX-induced DNA damage leads to cell death, and how AMH might be rescuing the DNA-damage response. To examine DOX-induced apoptosis, we performed a TUNEL assay across conditions (Figure S5). At 4h, 24h, and 1-week post-treatment we did not observe any signs of apoptosis in primordial oocytes, pre-granulosa cells, or granulosa cells of primary follicles; however, TUNEL staining increased in the granulosa cells of growing follicles of the secondary and antral stage. This was most evident with the increased number of TUNEL-positive granulosa cells at the 4h time points when comparing the control and DOX-treated ovaries (Figure S5). In contrast, AMH-DOX co-treatment significantly reduced the number of TUNEL-positive granulosa compared to DOX alone at the 4h and 1-week timepoints in antral follicles (Figure S5).

To understand how the DNA damage caused by DOX is resolved, we examined the accumulation of unresolved DNA damage foci by IHC using an antibody marking γ-H2AX, a marker of double-stranded DNA break (DSB). We observed an increase in γ-H2AX foci in the nucleus of primordial and primary oocytes in the DOX-treated group. Quantification of γ-H2AX positive follicles revealed that AMH cotreatment with Doxorubicin significantly reduced the accumulation of γ-H2AX in oocytes of primordial and primary follicles (Figure 5A). In secondary or larger follicles, γ-H2AX was detected in granulosa cells but not in oocytes, suggesting differential susceptibility of germ cells and granulosa cells to DOX-induced DNA damage, which varied by stage. To better understand how granulosa cells of growing follicles may be responding to DNA damage we quantified the ratio of positive granulosa cells per follicle and found a trend for an increased proportion of γ-H2AX positive granulosa cells in all follicle types following DOX treatment, reaching statistical significance in secondary follicles. Importantly, we also observed a trend for AMH co-treatment to reduce the proportion of γ-H2AX positive granulosa cells which was statistically significant in antral follicles (Figure 5A). These findings suggest DOX strongly induced DNA damage in oocytes of primordial and primary follicles and GC of larger secondary and antral growing follicles, while co-treatment with AMH resulted in a significant decrease in the accumulation of unresolved DSB across these follicle types and cells.

**Figure 5.**
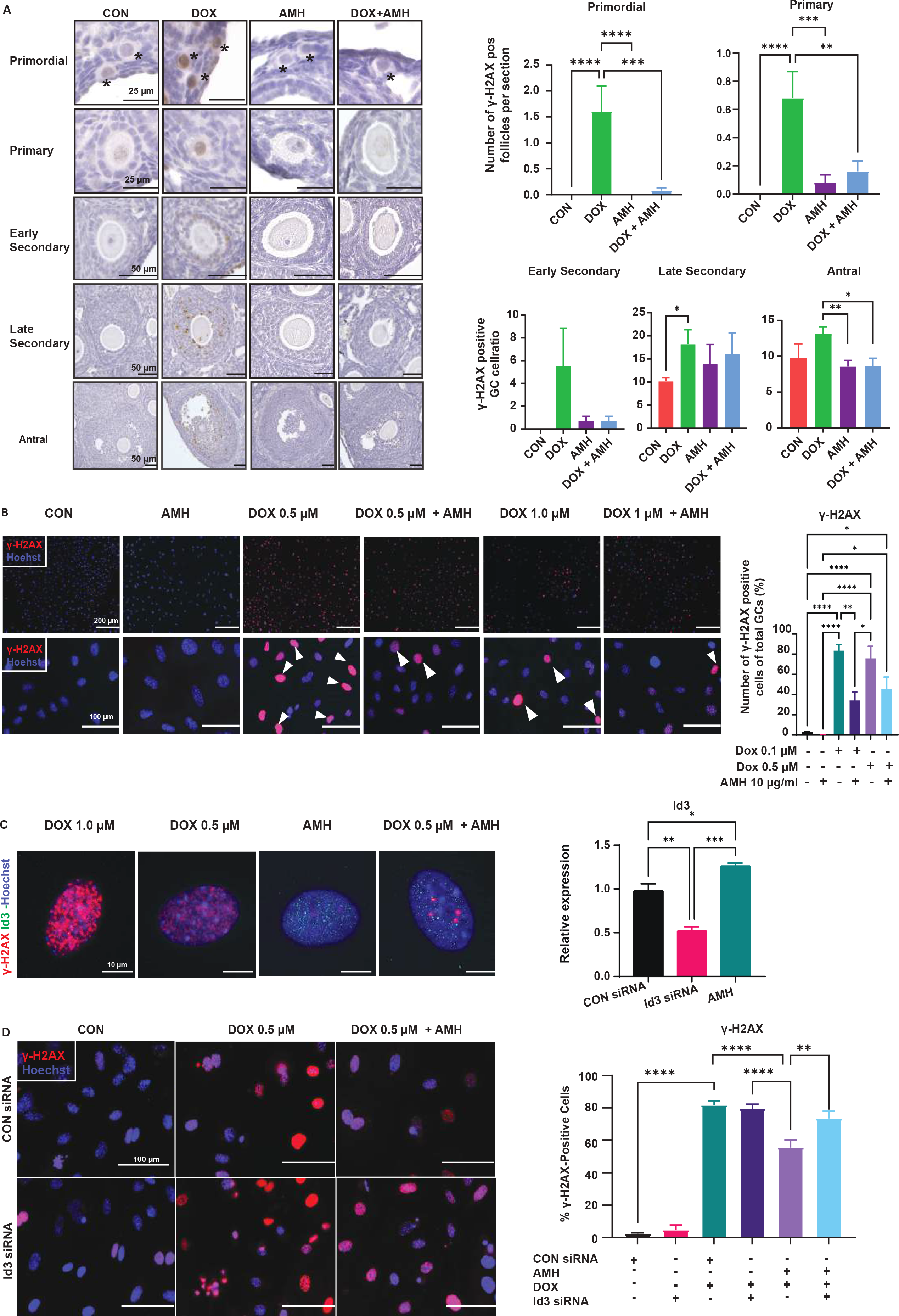
DOX and AMH effects on γ-H2AX accumulation. (A) Left panel: representative immunohistochemistry of γ-H2AX, an early marker of DNA damage in each treatment group, with black asterisks highlighting the primordial follicles. Right panel: Quantification of γ-H2AX-positive follicles. For primordial follicle and primary follicle >1 positive cell was considered a positive follicle. For secondary and larger follicles, the ratio between γ-H2AX positive granulosa cells and the total number of granulosa cells per follicle was calculated (n=30). Early secondary follicles were defined as surrounded by 2 layers of granulosa cells. Late secondary contained >2 layers of granulosa cells without antrum. Scale bars represent 25 µm for primordial and primary follicles, 50 µm for secondary and antral follicles. Data are presented as mean ± SEM; ∗p < 0.05, ∗∗p < 0.01, ∗∗∗p < 0.001. (B) Left panel: Representative micrograph of granulosa cells in vitro stained by immunofluorescence for γ-H2AX after treatment with different concentrations of DOX, with or without co-treatment of AMH (10 µg/ml). The bottom panel shows a higher magnification for each condition. Scale bars, 200 µm (top panel), 100 µm (bottom panel). Right panel: Quantification of γ-H2AX signal by image analysis with (B), mean number of γ-H2AX positive cells as a proportion of total granulosa cells. Data are presented as mean ± SEM; ∗p < 0.05, ∗∗p < 0.01, ∗∗∗p < 0.001. (C) Representative immunofluorescence stain of γ-H2AX and ID3 in granulosa cells. ID3 protein expression was evaluated after treatment with DOX (0.5 µM or 1 µM) and AMH (10 µg/ml). (B) Quantitative PCR of *Id3* with AMH treatment or siRNA knockdown. Data are presented as mean ± SEM; *p < 0.05, **p < 0.01, ***p < 0.001. (D) Representative micrograph of granulosa cells in vitro treated with an siRNA against *Id3*, or a control scrambled siRNA, with Dox (0.5 µM) and/or AMH (10 µg/ml) treatments, stained by immunofluorescence for γ-H2AX. Image quantification represents the percentage of γ-H2AX positive cells across conditions. Data are presented as mean ± SEM (N=5 experiments); *p < 0.05, **p < 0.01, ***p < 0.001.

We next sought to understand if the protective effect of AMH against was mediated directly on GCs or required other cell types of the follicle (oocyte, theca) to be present. We therefore isolated mitotically-active undifferentiated GCs from early-growing follicles collected from ovaries following diethylstilbestrol (DES) stimulation in prepubertal day 21 mice. We then treated these cells with either DOX only (0.5 μM and 1.0 μM for 48h) or with AMH (10 μg/ml for 76h including 24h pretreatment) or vehicle control in culture, and evaluated their ability to clear DNA damage by measuring γ-H2AX as a marker of unresolved DSBs. Per our in vivo results, GCs in culture also showed reduced γ-H2AX accumulation in DOX+rhAMH treated GCs compared to the DOX alone group, as indicated by the percentage of γ-H2AX positive cells (Figure 5B), suggesting a direct effect of AMH.

To understand how GCs recover from DOX damage we compared AMH+DOX cotreated- cells to DOX alone in a clonogenic assay. This allowed us to expose cells to DOX for a short period, and monitor their long-term recovery by the number and size of colonies, reflecting clonogenic cell survival, recovery, and proliferation. GCs (1.0×10^4^ / well) were seeded in 6 well plates and pre-treated with AMH for 24h followed by the addition of 0.1 µM or 0.5 µM of DOX for 48h; cells were then grown in drug-free medium for 10 days. Total colony size was quantified by crystal violet staining and absorbance, which was increased in the DOX+AMH group compared to DOX alone (DOX 0.1 µM vs DOX 0.1 µM + AMH 10 μg/ml, 1.69-fold increase, P< 0.001; DOX 0.5 µM vs DOX 0.5 µM + AMH 10 μg/ml, 1.42-fold increase, P< 0.05) (Figure S5B).

Interestingly, AMH pre-treatment alone also conferred a subsequent advantage in cell growth (albeit less pronounced) compared to control-PBS-treated cells, even in the absence of drugs (Figure 5S5B), which parallels our observation of rebound in theca cell progenitors following treatment (Figure S3E). These data suggest that co-treatment with AMH confers a survival and proliferative advantage to GCs after DOX exposure.

Finally, to determine if Id3 induction by AMH directly contributed to the decreased vulnerability of GCs to DOX, we used an siRNA to knock-down its expression in vitro. Indeed, increased ID3 protein expression in the nuclei of GCs after AMH treatment correlated with a reduction in γ-H2AX by immunofluorescent staining following treatment with 0.5 µM of DOX (Figure 5C). This AMH-dependent increase of *Id3* transcript was also evident at the transcriptional level, and *Id3* could be knocked-down using an siRNA (Figure 5C). As expected, treatment with 0.5 µM of DOX induced an accumulation of γ-H2AX staining by immunofluorescence and image analysis, which was significantly reduced by treatment with AMH; however, the γ-H2AX accumulation could be significantly rescued by knocking down Id3 (Figure 5D), suggesting AMH promotes resolution of DNA damage in part through upregulation of Id3.

## Discussion

In this study, we report the first single-cell atlas describing ovarian Doxorubicin fertotoxicity in prepubertal mice. We used this resource to elucidate the response of various ovarian cell types to DOX and/or AMH. We focused on this particular chemotherapy because our previous study suggested that gene therapy and recombinant AMH protein was most effective at protecting the ovarian reserve of mice treated with repeated dosing of DOX^11^.

The mechanism of action of DOX in the context of its use for chemotherapy is its generation of DNA damage, by acting as an intercalating agent that disrupts topoisomerases and causes DNA adducts, single-strand breaks, and double-strand breaks, particularly in actively dividing cancer cells^31^. However, these same types of DNA lesions can also occur in other benign cells, contributing to severe side effects including cardiotoxicity^32^ and fertotoxocity^11,33^. In the context of the ovary, genotoxic chemotherapies such as Doxorubicin are thought to contribute to accelerated ovarian aging by causing a loss of ovarian reserve through apoptosis of the highly susceptible primordial oocyte^34–36^, or by accelerating the rate of activation of primordial follicles, either in response to the injury itself^18^ or via loss of negative feedback paracrine inhibitors such as AMH, normally secreted by growing follicles, which are otherwise depleted by treatment^37^. Doxorubicin is also known to induce cellular senescence^38^, and may contribute to the deterioration of the ovarian stroma, and particularly the ovarian microvasculature, which could in turn adversely impact follicles^19^.

Herein, we identified pathways consistently activated by DOX across various cell types of the ovary and propose multiple potential protective mechanisms of AMH that may be cell-type specific and contribute to the preservation of ovarian health. The response to DNA-damage response includes recruitment of the repair machinery to the break site marked by γ-H2AX, signaling of the checkpoint kinases CHK1 and CHK2, and if unresolved, can lead to activation of p53 and in some cases its downstream target p21 (*Cdkn1a*), which leads to cell cycle arrest, senescence, and apoptosis ^31^. Likewise, in the ovary, we observed a consistent upregulation of *Cdkn1a* across multiple cell types. *Cdkn1a* is an important negative regulator of the cell cycle and transcriptional target of p53^39,40^, which has been previously reported to be induced by DOX in prepubertal mice^38^. Our findings suggest that DNA damage by DOX activates the p53 pathway, in turn inducing downstream effectors such as *Cdkn1a, Ccng1, Mdm2, Phlda3*, and ultimately resulting in the activation of the intrinsic apoptotic signaling pathway with upregulation of markers such as Bax. We found that these pathways were notably engaged in both the theca and preantral granulosa cells of the follicle. Importantly, markers such as Ccng1 and Mdm2 are components of a cell cycle checkpoint control axis that regulates proliferative cell competence, DNA fidelity, and survival^41^. The Ccng1/Mdm2/p53 axis has emerged as a strategic target for new precision molecular and genetic cancer therapies as well as chemo-sensitization^41^. While AMH has been proposed to contribute to the retention of the ovarian reserve following chemotherapy by preventing excessive activation of primordial follicles^11,14,42,43^, its potential salutary effects on growing follicles and other cell types of the ovary, and its involvement in the response to DNA-damage have not been thoroughly explored.

Unexpectedly, we found that AMH treatment may directly mitigate the genotoxic stress induced by DOX in growing follicles. Indeed, AMH treatment significantly diminished the expression of p53 class mediators normally induced by DOX in several mesenchymal and granulosa cell subtypes, including those of growing follicles. These results suggested that AMH could either suppress the DNA damage response and/or promote a faster resolution of DNA damage in ovaries. This was evident by the decrease of γ-H2AX-positive follicles in ovaries, and granulosa cells in vitro with AMH+DOX co-treament compared to DOX alone. Interestingly, γ- H2AX foci were more prevalent in oocytes of primordial follicles, compared to oocytes other than other types of follicles, whereas actively dividing granulosa cells of larger secondary and antral follicles accumulated more γ-H2AX compared to smaller follicles, suggesting different profiles of DOX follicular toxicity by stage and cell types. Surprisingly, AMH reduced the accumulation of γ-H2AX in the oocyte of primordial and primary follicles and in the granulosa cells of larger growing follicles. Since oocytes do not express AMHR2^8^, we speculate that any effects in germ cells are likely the result of AMH-driven juxtacrine signaling between oocyte and granulosa cells^8^. One way in which AMH may avert the accumulation of DNA damage in growing follicles is by reducing the susceptibility of cells to genotoxicity through inhibition of proliferation and/or by promoting DNA repair. To evaluate the contribution of these potential mechanisms, we took advantage of our scRNAseq dataset to build cell-state trajectory models using RNA velocity, and found that AMH stalled the differentiation of thecal progenitor cells into steroidogenic theca, resulting in a decreased expression of steroidogenic enzymes. These results are consistent with previous reports suggesting AMHR2 is expressed in theca cells and that AMH may inhibit steroidogenesis in part by inhibiting transcription of CYP17A1^44–47^. The stall in thecal progenitor differentiation was also accompanied by a reduction in proliferation which may also indirectly protect against genotoxic stress^48–51^. Notably, this hiatus in theca cell differentiation could be reversed, with the theca layer thickening one week after AMH treatment was discontinued. Protecting theca function could represent an important component of the oncofertility armamentarium, given the central role these cells play in maintaining steroid homeostasis, which is necessary for follicle growth and general endocrine health^52^.

In growing granulosa cells, we found a significant upregulation of *Wnt6* by DOX, particularly in preantral follicles. *Wnt6* is a critical determinant of pre-granulosa cell differentiation and subsequent oocyte maturation during primordial follicle activation^53^. We speculate that the upregulation of *Wnt6* might drive the overactivation of follicles, while its absence with AMH treatment could contribute to the suppression of follicle activation and growth. Indeed, following AMH treatment follicle progression from primary to secondary stage was also suppressed, as evidenced by an increased number of stalled primary follicles and fewer secondary follicles after one week of treatment. As exogenous AMH treatment is proposed to reduce primordial follicle activation, granulosa cell mitosis, and follicle-stimulating hormone-responsive follicle progression^8,54,55^, the capacity of AMH to stall follicles at an early stage may protect follicles by reducing primordial follicle activation and suppressing proliferation of granulosa cells of pre-antral follicles, which would otherwise render them susceptible to genotoxic stress. Thus, the manipulation of the WNT signaling pathway by AMH or other pharmacological means may offer a new potential target for therapeutic intervention.

Because many chemotherapies also damage non-proliferating cells, including oocytes of primordial follicles^9,11,19^, it had been unclear if AMH could protect such cells^11^. Surprisingly, our findings suggest that AMH may prevent the accumulation of DNA damage in both granulosa cells and oocytes. AMH induced a strong upregulation of *Id3*^8^, an important transcription factor that also plays a crucial role in DNA repair^30^, in both cell types. We observed a significantly reduced ability of AMH to rescue γ-H2AX foci accumulation in GCs when knocking down *Id3* in vitro. We therefore hypothesize that the upregulation of *Id3* by AMH may promote more effective DNA repair in cells affected by DOX and contribute to the enhanced survival of follicles. Future studies should investigate how DNA repair and follicle survival is coordinated between GC, oocyte, and theca cells, and aim to identify other drugs that ameliorate DNA damage response, such as Chek2 inhibitors^56^, particularly in oocytes, that could synergize with AMH.

In conclusion, this study provides new insights into how AMH may protect ovarian function and reveals multiple concurrent mechanisms of action, which combined, may mitigate some of the fertotoxicity of DOX. We speculate that some of these mechanisms of toxicity and rescue are likely to be implicated in other chemotherapies and in ovarian aging, given that DNA damage and follicle depletion are common. Furthermore, we chose to use prepubertal mice in this study to further the clinical development of AMH for the treatment of young cancer survivors, but more research is needed to ensure the efficacy and safety of AMH in humans.

## Limitations of the study

The dosage of DOX was chosen to facilitate the exploration of fertotoxic pathways and avoid mortality in mice and may not be directly translatable to clinical regimens used in pediatric cancer patients. The mechanisms of DOX toxicity may be dose-dependent and not broadly generalizable to other treatment regimens. Additionally, oocytes are poorly represented in our scRNAseq data, which reduced our ability to understand the consequences of treatments on these cells. Moreover, there is a broad need to understand more precisely the mechanistic conservation of fertotoxicity and AMH from mice to humans to ensure comparable safety and efficacy.

## Acknowledgments

We thank Philippe Godin for constructive feedback on the manuscript, and Caroline Coletti and Isabel Davis for administrative help. This study was funded by an NIH/NICHD grant (1R01HD102014) to D.P. The illustrations were created with BioRender.com.

## Author Contributions

Conceptualization, NMP.N., E.C., and D.P..; methodology, NMP.N., E.C., and D.P.; formal analysis, NMP.N., E.C., M-C.M., and D.P.; investigation, NMP.N., E.C., M.C., N.S., A.K., N.N., C.C., P.M, A.M., J.C., M-C.M., T.D.; data curation: NMP.N. and D.G.; writing – original draft; NMP.N., E.C., and D.P.; visualization, NMP.N.; writing – review & editing, NMP.N., P.K.D. and D.P.; supervision, NMP.N., E.C., M-C.M., and D.P.; funding acquisition, D.P.

## Declaration of interests

### Related patent filings

PCT/US2017/066346 - WO2018112168 - MULLERIAN INHIBITING SUBSTANCE (MIS) PROTEINS FOR OVARIAN AND UTERINE ONCOPROTECTION, AND OVARIAN RESERVE AND UTERINE PRESERVATION (D.P. and P.K.D.)

### Advisory and consulting position

Scientific advisors and co-founders of Oviva Therapeutics (D.P. and P.K.D).

## STAR Methods

### KEY RESOURCES TABLE

| REAGENT or RESOURCE | SOURCE | IDENTIFIER |
| --- | --- | --- |
| Antibodies |  |  |
| pH2AX | Cell Signaling Technology | Cat# 2577;<br>RRID:AB_2118010 |
| Ki67 | Thermo Fisher Scientific | Cat# 14-5698-80 |
| Chemicals, peptides, and recombinant proteins |  |  |
| LR-AMH | D. Pepin at al.<sup>57</sup> | N/A |
| Collagenase | Sigma Aldrich | Cat# 10103578001 |
| Papain | Worthington | Cat# LK003178 |
| Critical commercial assays |  |  |
| Chromium Next GEM Single Cell 3' Reagent Kits v3.1 (Dual Index) | 10× Genomics | PN-1000268 |
| Dual Index Kit TT Set A, 96 rxns | 10× Genomics | PN-1000215 |
| RNAscope™ 2.5 HD Assay - RED | Advanced Cell Diagnostics, ACD | Cat# 322350 |
| RNAscope Multiplex Fluorescent Reagent Kit v2 assay | Advanced Cell Diagnostics, ACD | Cat# 323110 |
| Deposited data |  |  |
| scRNA sequencing raw and processed data | This paper | GEO: GSE266600<br>Token:<br>yrghiceifxkxvot |
| Original codes | This paper | DOI:<br>10.5281/zenodo.11187609 |
| Experimental models: Organisms/strains |  |  |
| WT mice | Jackson Laboratory | RRID:IMSR_JAX:000664 |
| Oligonucleotides |  |  |
| RNAscope probe Mm-Cdkn1a-C2 | Advanced Cell Diagnostics, ACD | 408551-C2 |
| RNAscope probe Mm-Mdm2-C2 | Advanced Cell Diagnostics, ACD | 447641-C2 |
| RNAscope probe Mm-Trp53inp1-C1 | Advanced Cell Diagnostics, ACD | 1161531-C1 |
| RNAscope probe Mm-Ccng1-C1 | Advanced Cell Diagnostics, ACD | 433221 |
| RNAscope probe Mm-Bax | Advanced Cell Diagnostics, ACD | 463501 |
| RNAscope probe Mm-Phlda3 | Advanced Cell Diagnostics, ACD | 476051 |
| RNAscope probe Mm-Prss35-C2 | Advanced Cell Diagnostics, ACD | 492611-C2 |
| RNAscope probe Mm-Cyp17a1 | Advanced Cell Diagnostics, ACD | 522611 |
| RNAscope probe Mm-Cyp11a1 | Advanced Cell Diagnostics, ACD | 809181 |
| RNAScope probe Mm-Clec3b | Advanced Cell Diagnostics, ACD | 539561 |
| RNAScope probe Mm-Pgr-C2 | Advanced Cell Diagnostics, ACD | 318921-C2 |
| RNAScope probe Mm-Id3 | Advanced Cell Diagnostics, ACD | 445881 |
| Software and algorithms |  |  |
| Seurat | Butler et al.<sup>58</sup> | https://cran.r-project.org/web/packages/Seurat/index.html |
| ClusterProfiler | Yu et al.<sup>59</sup> | https://yulab-smu.top/biomedical-knowledge-mining-book/universal-api.html |
| GSEA | Subramanian et al.<sup>60</sup> | http://www.broad.mit.edu/gsea/ |
| Velocyto | La Manno et al.<sup>61</sup> | https://velocyto.org/velocyto.py/index.html |
| CellChatDB | Jin et al.<sup>29</sup> | www.cellchat.org/cellchatdb |
| ImageJ | Schneider et al.<sup>62</sup> | https://imagej.net/ij/download.html |

### RESOURCE AVAILABILITY

#### Lead contact

Further information and requests should be directed to the lead contact, David Pépin

### Materials availability

This study did not generate new unique reagents.

#### Data and code availability

- The single-cell RNA-seq data generated in this study have been deposited at Gene Expression Omnibus with the accession number GEO: GSE266600. Token: yrghiceifxkxvot.
- Microscopy data reported in this paper will be shared by the lead contact upon request.
- All original code has been deposited at Zenodo and is publicly available as of the date of publication. DOIs are listed in the key resources table.
- Any additional information required to reanalyze the data reported in this paper is available from the lead contact upon request.

### EXPERIMENTAL MODEL AND STUDY PARTICIPANT DETAILS

#### Mouse models

This study was conducted under an experimental protocol (2019N000068) approved by the Massachusetts General Hospital Institutional Animal Care and Use Committee. The mice of strain C57BL/6J were purchased from the Jackson Laboratory. Female mice, 25 days postnatal, were treated with rhAMH through intraperitoneal injections twice daily. The DOX group received two intraperitoneal injections at a one-week interval, starting on postnatal day 26, with a dosage of 3 mg/kg. Mice were euthanized at 4 hours, 24 hours, and 1 week after the last DOX dose by inhalation of CO2, and then tissue and blood samples were collected. Additionally, on postnatal day 25 (day 0), the mice received either rhAMH or saline as a vehicle control.

#### Isolation and culture of mouse primary granulosa cells

21-day-old female C57BL6 mice were treated with 100μg diethylstilbesterol (DES) dissolved in Corn Oil (1mg/1ml) via subcutaneous injection. After 48-h treatment, the mice were killed and the ovaries were removed aseptically. After ovaries were pooled and washed in the isolating medium, GCs were collected immediately by puncturing the pre-antral or small antral follicles from each ovary with a 25-gauge needle in a Falcon dish (Greiner, Frickenhausen, Germany) and the cell suspension was filtered with a 40μm cell strainer (Corning, #431750). The GCs were resuspended and diluted in McCoy’s 5A supplemented with 5% Fetal Bovine Serum, 1% penicillin with streptomycin sulfate, and 1% Insulin-Transferrin-Selenium. Cells (10^5^ cells/ml) were then cultured at 37℃ in a humidified atmosphere of 95% O2 and 5% CO2 for further study.

### METHOD DETAILS

#### Histological follicle counts

Ovaries were fixed in 4% paraformaldehyde overnight at 4℃ and then processed through a series of graded alcohols and xylenes baths for 24 hours prior to paraffin embedding. Embedded tissues were sectioned serially (5μm thickness) and then placed on Superfrost Plus staining slides (Thermo Fisher Scientific, Waltham, MA, USA). For follicle counting with an accurate estimation of the size of primordial follicles, every tenth serially sectioned representative slides (sections per group=25) were immunostained with DEAD-Box Helicase 4 (DDX4) (1:1000, ab27591, Abcam, USA). Hematoxylin (Mayer’s; Dako, Carpinteria, CA) was used for nuclear counterstaining. The sections were mounted with Cytoseal on microscope slides (Fisher Scientific), and follicle counts were performed. The follicle stage was classified according to previously accepted definitions^63^. A primordial follicle (PMF) was defined as an oocyte surrounded by a single layer of flattened squamous pre-granulosa cells. A primary follicle (PF) was defined as an oocyte surrounded by a single layer of cuboidal granulosa cells (GCs). Occasionally, follicles appeared as an intermediate stage between the primordial and primary stages, with both cuboidal and squamous GCs. If the cuboidal cells predominated, the follicle was classified as a PF, they were otherwise classified as PMF. Secondary follicles (SF) had two or more layers of cuboidal GCs with no visible antrum. Antral follicles (AF) were defined as follicles with an antral space containing follicular fluid. Atretic follicle (ATF) were the follicles with more than 50% of apoptotic GCs and degenerative oocytes. After the quantification of follicles, the mean number of follicles in each stage per section was calculated.

#### Generation of single-cell suspension for scRNAseq

One ovary from each animal was collected in cold PBS, and the pooled ovaries from each group (N = 5 per group) were cut into small pieces and placed in 2 mL of DMEM (10% FBS, 1% penicillin/streptomycin), supplemented with 21 U/mL of Papain (Worthington, LK003178) and 1 mg/mL of collagenase A (Sigma Aldrich, 10103578001), and incubated for 30 minutes at 37°C. After digestion, the cells were subjected to further mechanical digestion using the gentleMACS™ Dissociator (Miltenyi Biotec). Single cells were isolated by straining the digested tissue suspension through a 70-micron filter. The cells were then quantified to assess viable cell concentration and 10,000 cells from each group were collected for library preparation. The other ovary from each animal was placed in 10% neutral buffered formalin at room temperature overnight before being processed using an automated tissue processor for histological analyses.

#### Library preparation, sequencing, and data processing

Samples were processed for scRNAseq according to the manufacturer’s instructions using the Chromium Next GEM Single Cell 3ʹ Reagent Kits v3.1 (Dual Index) (10× Genomics) and Chromium Controller (10× Genomics). Single-cell libraries were sequenced using the NovaSeq S4 (Illumina) by the Bauer Core Facility at Harvard University.

Data analysis was performed using CellRanger (version 7.0.0) with standard parameters. The samples were aligned against the mouse reference transcriptome mm10-2020-A and analyzed using Seurat ^58^ (R version 4.2.0, Seurat version 4.1.0). A standard pre-processing workflow was applied independently to each sample. Samples were filtered based on a mitochondrial percentage under 20% and unique feature counts between 200 and 10,000. After normalization, samples were merged using the ’merge’ function in Seurat. Conventional visualization and clustering methods were applied, and cluster-specific markers were identified with the ‘FindAllMarkers’ function in Seurat.

#### Differentially Expressed Genes and Pathway Analyses between Different Treatments

Significantly differentially expressed genes between treatments were identified using the Seurat ‘FindMarkers’ function. Volcano plots were generated using ‘EnhancedVolcano’ functions. Differentially expressed genes with adjusted p-values less than 0.05 were used as input for gene ontology enrichment analysis by the clusterProfiler package ^59^. The biological process subontology was chosen for this analysis. The Enrichplot package was used for the visualization of Functional Enrichment Results ^64^

All GSEA analyses were performed using GSEA software^60^ with an unranked method. Counts were extracted from the Seurat object using the ‘GetAssayData’ function as the expression dataset input (.txt). Phenotype label files (.cls) were created in Microsoft Excel. GSEA was performed with the following parameters: number of permutations: 1000, no collapse to gene symbols, permutation type: gene_set, gene set size 5-500. Either Hallmark signature or M2 gene sets (curated studies) were used.

#### Cellular differentiation analysis

The analysis was performed as described by La Manno et al.^61^, using the Velocyto package to generate unspliced and spliced counts and employing scVelo for RNA velocity analysis. Initially, a matrix for spliced and unspliced transcripts was constructed using the ’velocyto run10x’ command line tool with mouse reference transcriptome refdata-gex-mm10-2020-A downloaded from 10X Genomics. This generated properly formatted loom files from each BAM file. Seurat UMAP coordinates were extracted in R to be integrated into the RNA velocity analysis. Subsequently, loom files from all four groups were concatenated. Next, scVelo was run to compute RNA velocity. First, the data were preprocessed with filtering and normalization applied to the spliced and unspliced counts. RNA velocity was then computed using the steady-state model (stochastic option) via scv.tl.velocity(). Finally, RNA velocity was visualized using scv.pl.velocity_embedding_stream().

#### Cell-cell interaction analysis

Cell-cell communication calculation and analysis was performed using CellChat ^29^, an R-based analysis tool. Cellchat object was created using the ‘createCellChat’ function, with the Seurat object of each condition (CON or DOX) as input. The ‘CellChatDB.mouse’ database was used as the ligand-receptor interaction database. Each group was independently preprocessed and analyzed to infer the cell-cell communication network using the ‘computeCommunProb’ function. The inferred communication network for specific signaling pathways of interest (e.g., AMH) was visualized using ‘netVisual_aggregate’. To identify differences between groups, CellChat objects were merged using ‘mergeCellChat’. The number of interactions and interaction strength among different cell populations were compared using ‘netVisual_diffInteraction’. Specific signaling changes in preantral granulosa cells between the DOX and CON groups were identified using ‘netAnalysis_signalingChanges_scatter’, specifying ‘idents.use’ as the preantral GC cluster.

#### Immunohistochemistry

Paraffin sections were deparaffinized in xylene followed by rehydration through a series of ethanol. For antigen retrieval, sections were treated for 40 minutes at 95-100 °C with 0.01M sodium citrate buffer (pH 6.0). Sections were then washed with TBS containing 0.1% Tween 20 (TBST) and incubated in 0.3% hydrogen peroxide in distilled water for 10 minutes. The slides were then washed with TBS and blocking solution (2% BSA in TBST) was applied for 1 hour at RT. The sections were incubated with the antibody against pH2AX (1:500, cell signaling #2577), Ki67 (1:200, Thermo Fisher Scientific # **14-5698-80)** for 18 hours at 4°C, followed by incubation with diluted horseradish peroxidase-conjugated goat anti-Rabbit IgG antibody (1:400, Cell Signaling #7074S), HRP-conjugated donkey anti-Rat IgG antibody (1:500, Jackson laboratory #712-035-153) for 2 hours at RT. The slides were washed with TBST and placed in distilled water. 3,3’-Diaminobenzidine (DAB) and chromogen substrate solution (Dako, Carpinteria, CA, USA #K3468) was added dropwise onto the tissues according to the manufacturer’s protocol. The sections were counterstained with hematoxylin and dehydrated with ethanol and xylene, then mounted with coverslips using cytoceal 60 mountant (Fisher Scientific, US # 831016).

#### RNA in situ hybridization

In situ hybridizations were performed using ACDBio kits as per manufacturer’s protocol, and as previously described^12^. Briefly, RNAish was developed using RNAscope Mutiplex Fluorescent Reagent Kit v2 assay or the RNAscope 2.5 HD Reagent Kit (RED) (ACD Bio). Following deparaffinization in xylene, dehydration, peroxidase blocking, and heat-induced epitope retrieval by the target retrieval and protease plus reagents (ACD bio), tissue sections were hybridized with probes for the target genes in the HybEZ hybridization oven (ACD Bio). The slides were then processed for standard signal amplification steps and chromogen development.

#### TUNEL and Immunofluorescence

Cell death (TUNEL-assay) was detected using an in situ cell death detection kit (Click-iT Plus TUNEL Assay with Alexa 488, C10617, Invitrogen) according to the manufacturer’s instructions. DNase I-treated sections were used as a positive control for DNA breakdown in the follicle nuclei. The nuclei were stained with Hoechst 33342.

For Immunofluorescence, cells were plated in 6 well plate with gelatin coated coverslip. When cells reached desired density, culture media was removed and cells were washed with PBS, fixed with 4% PFA and permeabilized with 0.1 % TX-100/PBS. Subsequently, cells were blocked with 1% BSA/PBS for 45 minutes and incubated at 4°C overnight with primary antibodies, followed by 1h with secondary antibodies, all diluted in blocking buffer. After nuclear staining with Hoechst, coverslips were washed and mounted with Vectashield® antifade mounting medium. The primary antibodies used were LRH-1 (1:200, R&D system, PP-H2325), pH2AX (1:500, cell signaling #2577), Id3 (1:100, Santa Cruz, sc-56712), and for secondary antibodies, Alexa Fluor 594 donkey anti-rabbit IgG (1:200, # A-21207, Invitrogen) and Alexa Fluor 488 donkey anti-mouse IgG (1:200, ab150105, Abcam) was used.

All images were collected and analyzed using Keyence BZ-X800 Digital Microscope and software (Keyence Corporation of America). The number of positive cells in the ovary was counted using Image J software (NIH).

#### Clonogenic assay

For the clonogenic assay, primary GCs (10^4^ cells/well) were plated in six-well plates before treatment. Cells were treated with either Dox alone (0.1μM and 0.5μM for 48h) or with rhAMH (10 μg/ml for 76h including 24h pretreatment). Then, the medium was changed and cells were cultured for additional 10 days in a drug free medium. Subsequently, cell colonies were fixed and stained with a solution of 80% crystal violet and 20% methanol. Colonies were then photographed. Then the cells were washed (once with PBS) and 30% v/v acetic acid was added to induce a complete dissolution of the crystal violet. Absorbance was recorded at 595-600 nm in the VICTOR™ Multilabel Plate Reader (PerkinElmer, Waltham, MA, USA). Three different experiments were performed.

#### Id3 knockdown with siRNA

Twenty-five days old immature mice were superstimulated with a single subcutaneous injection of Diethylstilbestrol (100ug/mouse dissolved in Corn Oil, 1mg/1ml) to induce follicular growth and their ovaries were harvested 48h later as previously described (PMID: 21999365). Follicles were manually punctured with a 25-gauge needle, cells filtered (40um filter) and plated in 6 wells plate (0.5x106 cells/well) in supplemented McCoy5A culture medium (5% FBS/1% PenStrep/1%ITS) before being cultured at 37℃ in a humidified atmosphere of 95% O2 and 5% CO2 for three days before treatment. The cells were transfected with 50nM of siRNA targeting ID3 (s68012, s68013 ; Silencer® Select Pre-Designed siRNA; cat#4390771) or 50nM scramble non-targeting control siRNA (Silencer® Select Negative Control #1 ; cat#4390843) using lipofectamine RNAiMAX (Invitrogen) according to the manufacturer’s instructions. The cells were incubated in the transfection mixture for 5 hours. Then, they were further treated with Dox only (0.5μM for 24h) or Dox (0.5μM for 24h) with rhAMH (10 μg/ml for 48h including 24h pretreatment). The efficiency of the knock- down was validated by q-PCR.

Following fixation in 4%PFA for 10min, granulosa cells were permeabilized 1%TritonX, washed with 0.1% PBST and blocked in 2.5%BSA/PBST before being incubated overnight with γ-H2AX (1:200, cell signaling #2577) and Id3 antibody (1:100, Santa Cruz, sc-56712). The granulosa cells were then rinsed with PBST and incubated for 1h with secondary antibody Alexa Fluor 594 donkey anti-rabbit IgG (1:200, # A-21207, Invitrogen) and Alexa Fluor 488 donkey anti-mouse IgG (1:200, ab150105, Abcam) before being counterstained with Hoechst.

Fluorescent images were quantified in ImageJ to calculate the percentage of positive cells. Total cell count and regions of interest (ROIs) representing each cell were obtained from the blue channel by binary thresholding and the “Analyze Particles” function. ROIs were then overlayed on the red channel to obtain mean grey value per cell. The number of positive cells was determined using a set intensity threshold across conditions and divided by total cell count to obtain positive cell percentage. Data for each plate was averaged. One way ANOVA was performed using GraphPad Prism to evaluate significance.

### QUANTIFICATION AND STATISTICAL ANALYSIS

Statistical evaluation was conducted using GraphPad Prism 10.2.1, using an independent sample ANOVA test. p ≤ 0.05 was considered as significant. The exact value of biological replicates/animals (n) can be found in the figure legends and results section.

For the quantification of in-situ hybridization assay, color deconvolution was performed using the Fiji image processing package to separate the red dot signals from hematoxylin staining. The images showcasing red staining were then converted to 8-bit grayscale, and the threshold settings were manually adjusted to accurately delineate the areas with positive staining. The ’Area fraction’ option was selected to calculate the percentage of positive staining within the manually designated areas, such as the theca or granulosa cells at different stages. For quantification analysis, three random areas of interest (from each category: theca, stroma, or preantral granulosa) were selected for analysis on each slide. This approach was consistently applied across all slides, with one slide representing each mouse and each experimental group consisting of 3 to 5 mice. The same criteria were used for quantifying all markers, except for Bax. For Bax, due to the sections having fewer preantral follicles compared to sections for other markers, the positive percentage measurements of the selected areas were first averaged for each slide. This preliminary step was completed prior to conducting the statistical analysis. For the in-situ hybridization assay using RNAscope Multiplex Fluorescent Reagent Kit, the normalized mean intensity was measured for each slide.

## Supplemental figure titles and legends

**Figure S1:**
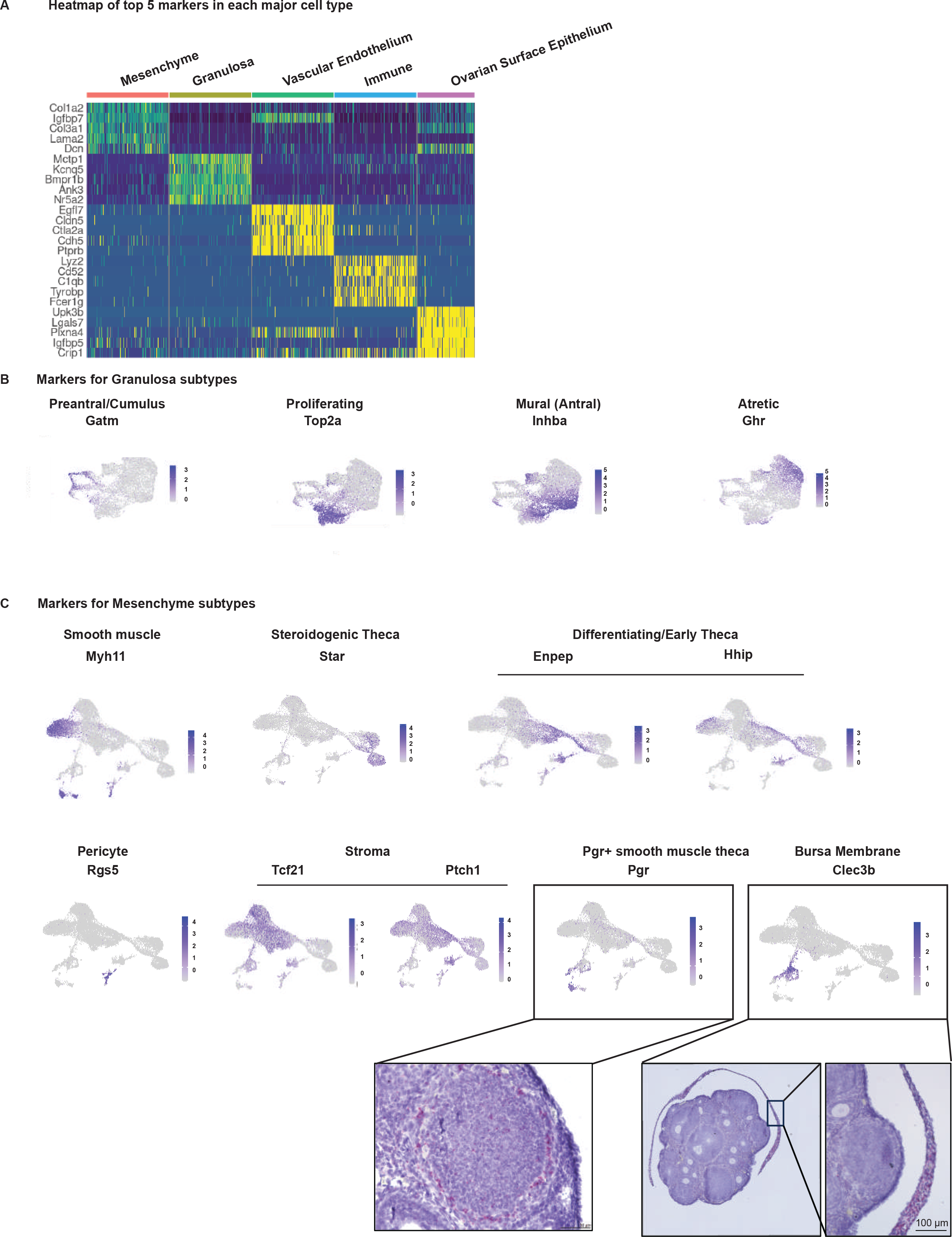
**Single-cell RNA sequencing analysis of mouse ovaries treated with DOX and AMH.** (A) Heatmap of the top 5 markers of each cluster by fold change. (B) Featureplots of selected markers for granulosa cell subtypes. (C) Featureplots of selected markers for mesenchymal cell subtypes.

**Figure S2.**
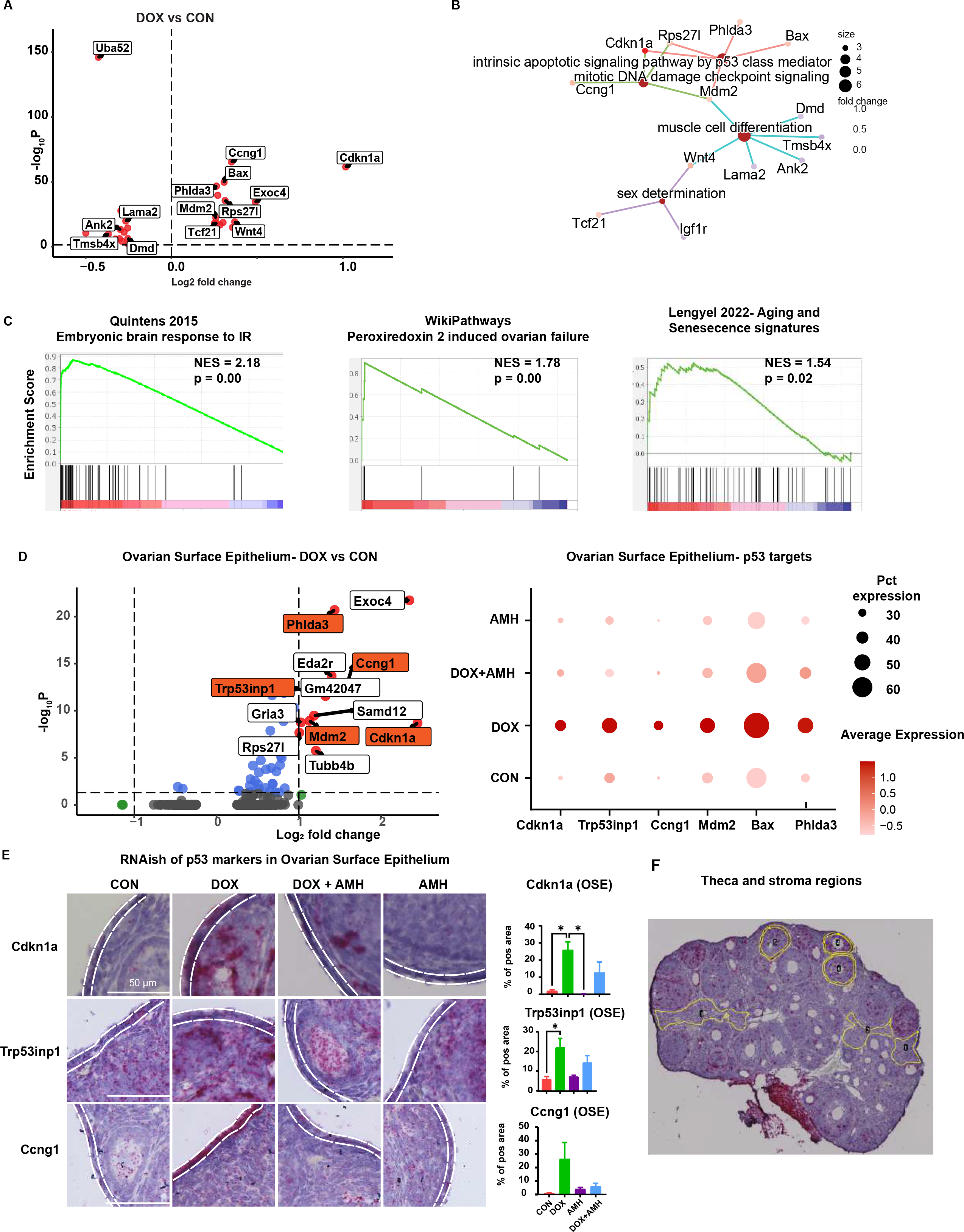
Effects of DOX on ovarian mesenchymal cells. (A) Volcano plot of DGE analysis comparing DOX-treated and control samples across all mesenchymal cells. (B) Gene Network featuring selected enriched pathways. (C) GSEA using the M2-mouse collection with observed enrichment of genes in response to IR, ovarian failure pathway, or ovarian aging when compared with the Lengyel et al. study (last panel). (D) Left panel: Volcano plot of DGE between AMH-treated and control in ovarian surface epithelium. Right panel: Dotplot of markers in p53 targets pathway activated by DOX in Ovarian surface Epithelium. (E) Representative micrograph of ovarian sections stained by RNAish for selected p53 markers in ovarian surface epithelium, with image quantification. Scale bars, 50 µm. Data are presented as mean ± SEM; ∗p < 0.05, ∗∗p < 0.01, ∗∗∗p < 0.001. (F) Representative micrograph of a low magnification ovarian section stained for *Cdkn1a* by RNAish for quantification in mesenchyme cells. For the theca layers, 3 follicles were selected per slide. For the stroma areas, 3 random areas were selected per slide.

**Figure S3.**
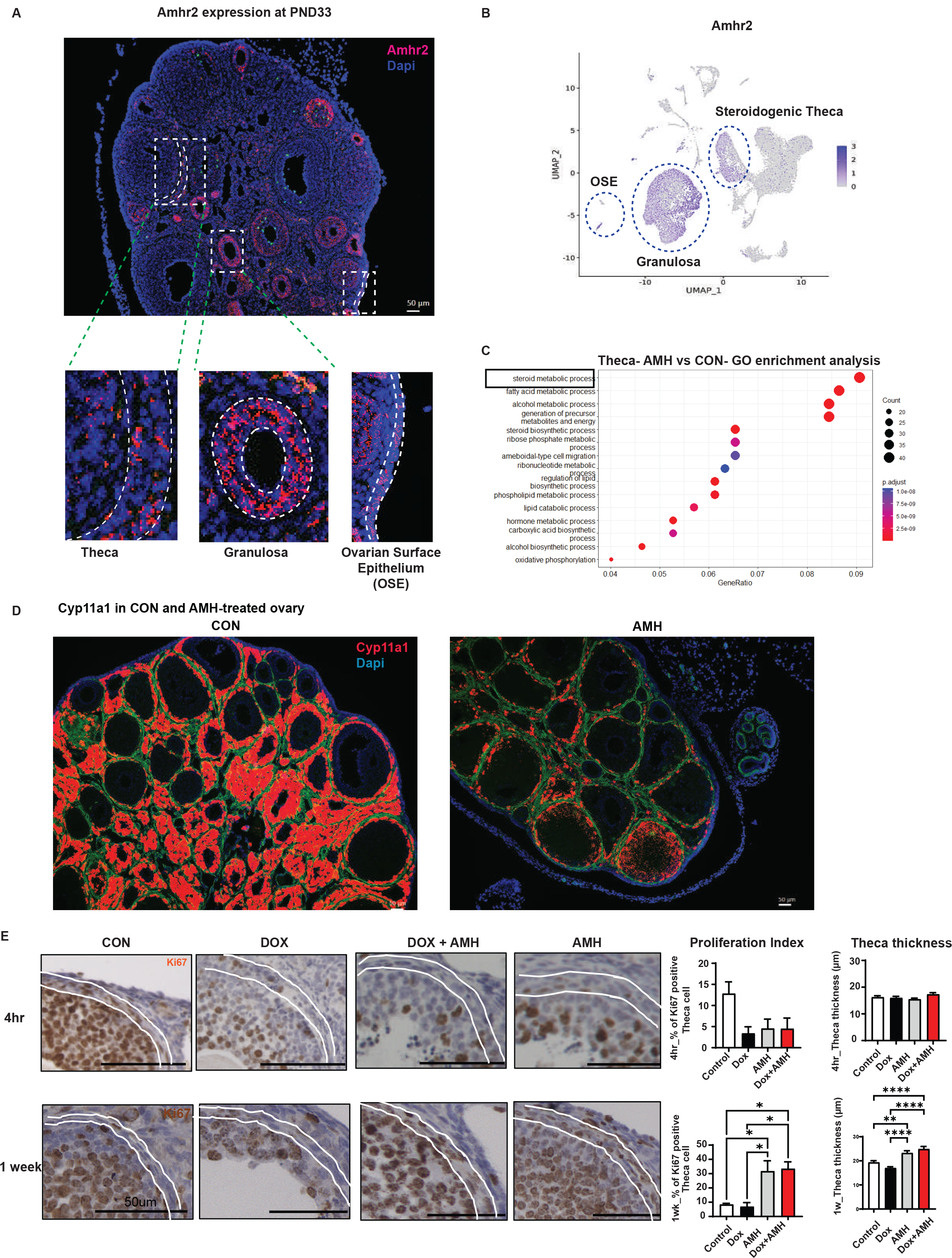
AMH impact on theca cells. (A) Representative micrograph of a section stained by fluorescent RNAish for *Amhr2* in the ovary at PND33. (B) Featureplot of *Amhr2* expression in different cell types. (C) GO analysis of AMH vs CON in theca revealing steroid metabolic process as the most enriched pathway. (D) Representative micrograph of a section stained by fluorescent RNAish for *Cyp11a1* in control and AMH-treated ovary. (E) Representative micrograph of ovarian section stained by IHC for Ki67 in ovaries at 4 hours and 24 hours after stopping AMH treatment, with proliferative index calculated by the percent of Ki67 positive cells compared to the total number of theca cells. The diameter of the theca layer thickness in four experimental groups was also quantified.

**Figure S4.**
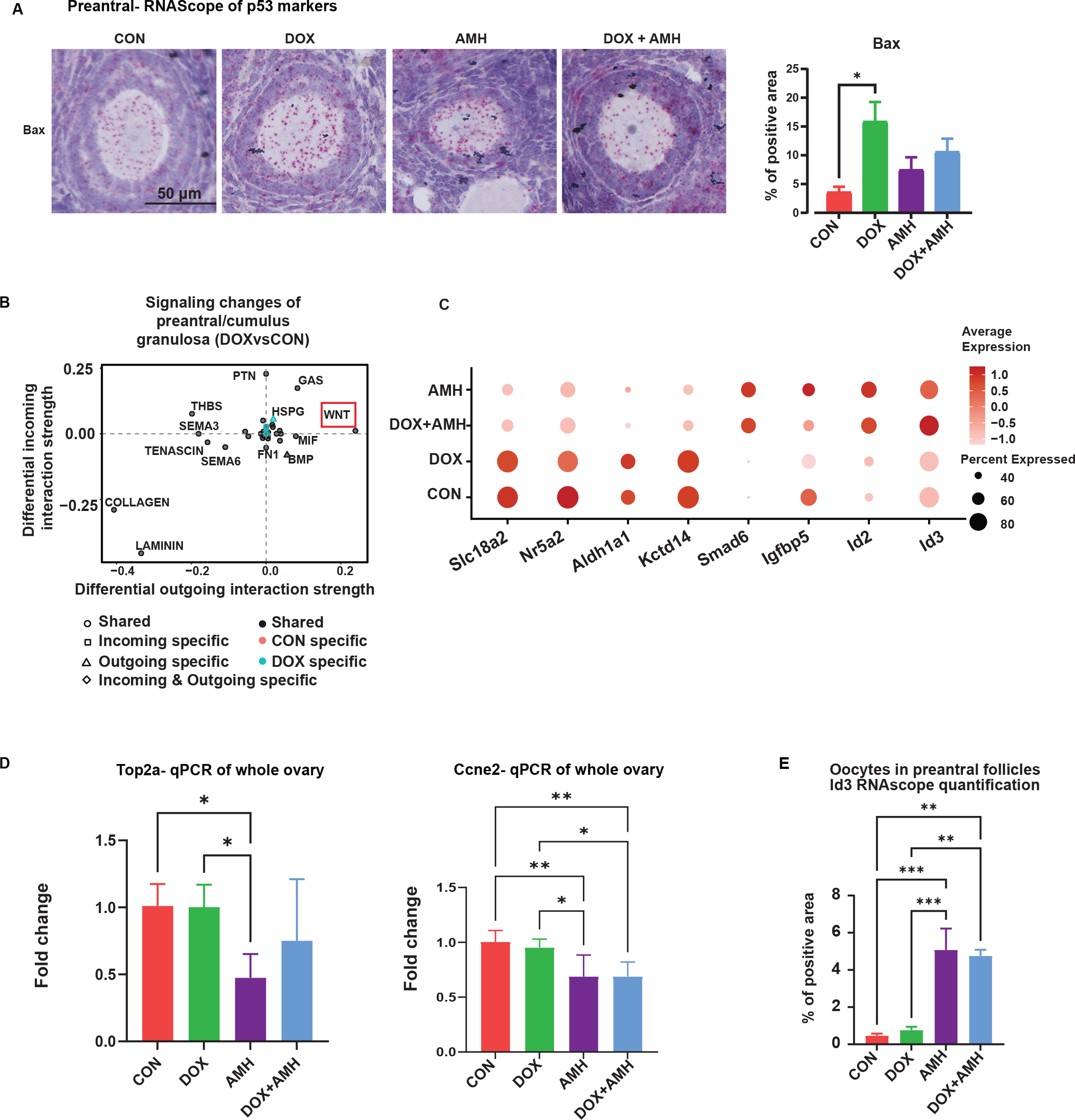
DOX impact on preantral granulosa cells. (A) Representative micrograph of an ovarian section stained by RNAish of apoptosis marker, Bax, in preantral, with quantification. Scale bars, 50 µm. Data are presented as mean ± SEM; ∗p < 0.05, ∗∗p < 0.01, ∗∗∗p < 0.001. (B) CellChat analysis of signaling pathways in preantral GC in ovaries treated with DOX compared to CON. The x and y axes represent differential outgoing and incoming interaction strengths, respectively. Positive and negative values indicate increased and decreased signaling, respectively, in ovaries treated with DOX compared to control ovaries. (C) Dotplot of specific gene signatures regulated by AMH treatment. (D) qPCR pf proliferation markers, *Top2a* and *Ccne2*, in whole ovaries across different conditions. (E) Image analysis of RNAish quantification of *Id3* in oocytes of preantral follicles. Data are presented as mean ± SEM; ∗p < 0.05, ∗∗p < 0.01, ∗∗∗p < 0.001.

**Figure S5.**
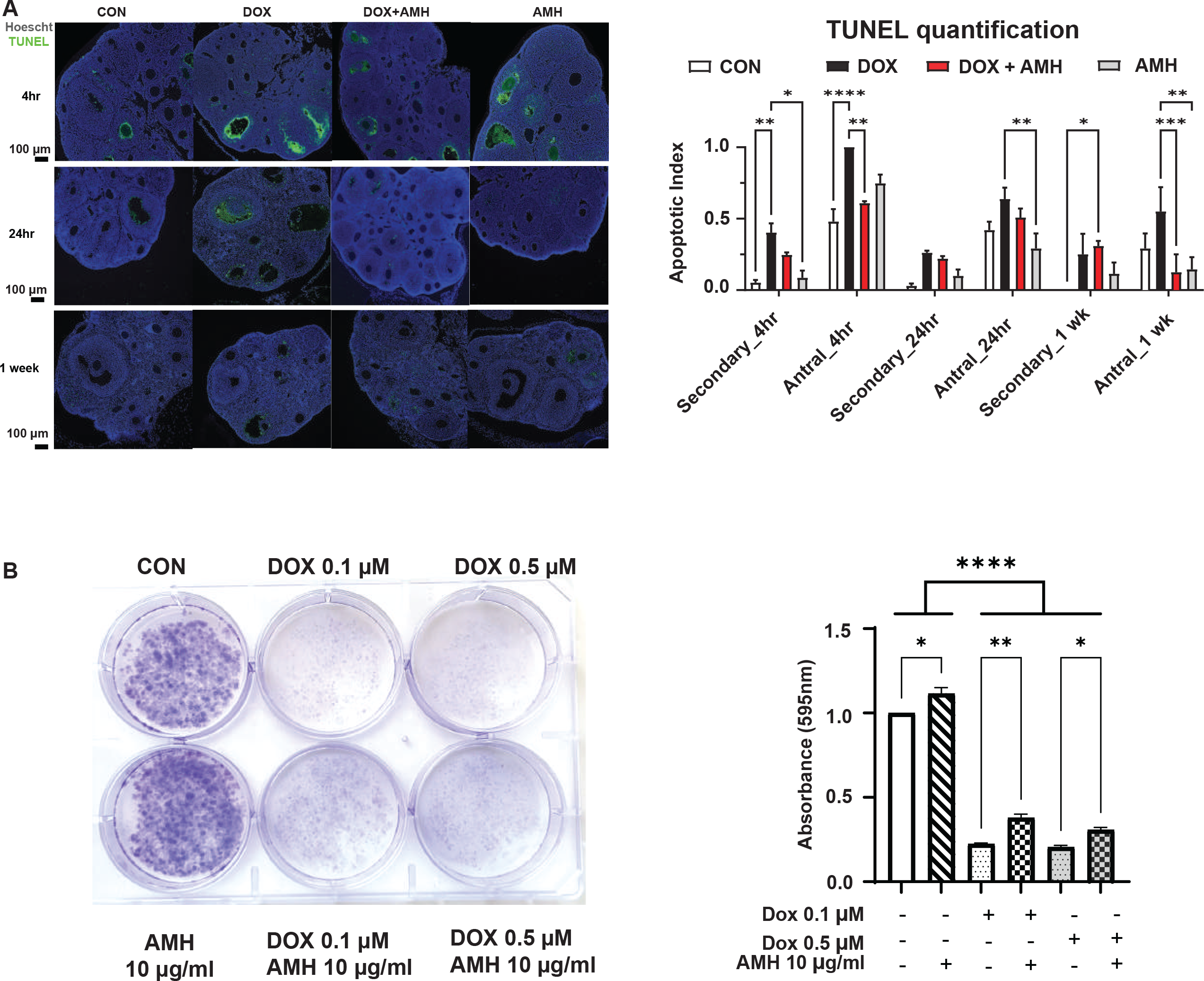
Impact of DOX and AMH on cellular apoptosis and survival. (A) Representative micrograph of an ovarian section stained by TUNEL with evidence of staining in secondary and antral follicles. Follicle with > 10 TUNEL-positive cells was considered as positive. The apoptotic index was calculated as the fraction of TUNEL positive follicles/total follicles for each follicular developmental stage. Data are presented as mean ± SEM; ∗p < 0.05, ∗∗p < 0.01, ∗∗∗p < 0.001. (B) Picture of a representative clonogenic assay performed in a six-well plate with primary immature granulosa cells. Cells were treated with DOX 0.1-0.5μM for 48h after a 24h pretreatment with AMH (N=3 experiments). The medium was changed and colonies were cultured for an additional 10 days in drug-free medium. Crystal violet staining was quantified by absorbance. Data are presented as mean ± SEM; ∗p < 0.05, ∗∗p < 0.01, ∗∗∗p < 0.001.

## Supplemental table titles

Table S1. Differential Gene Expression Analysis of Mesenchyme Clusters: Markers Differentiating DOX-Treated Ovaries from Controls. Related to Figure 2.

Table S2. Enriched Pathways in Mesenchyme Clusters Identified by Gene Ontology Enrichment Analysis: Comparison Between DOX-Treated Ovaries and Controls. Related to Figure 2.

Table S3. Enriched Pathways in the Proliferating Mesenchyme Cluster Identified by GSEA: Comparison Between DOX-Treated Ovaries and Controls. Related to Figure 2.

Table S4. Enriched Pathways in the Preantral Granulosa Cluster Identified by GSEA: Comparison Between DOX-Treated Ovaries and Controls. Related to Figure 4.

